# Optical Control of Membrane Fluidity Modulates Protein Secretion

**DOI:** 10.1101/2022.02.14.480333

**Authors:** Noemi Jiménez-Rojo, Suihan Feng, Johannes Morstein, Stefanie D. Pritzl, Takeshi Harayama, Antonino Asaro, Nynke A. Vepřek, Christopher J. Arp, Martin Reynders, Alexander J. E. Novak, Evgeny Kanshin, Beatrix Ueberheide, Theobald Lohmüller, Howard Riezman, Dirk Trauner

## Abstract

The lipid composition of cellular membranes is dynamic and undergoes remodelling affecting biophysical properties, such as membrane fluidity, which are critical to biological function. Here, we introduce an optical approach to manipulate membrane fluidity based on exogenous synthetic fatty acid with an azobenzene photoswitch, termed **FAAzo4**. Cells rapidly incorporate **FAAzo4** into phosphatidylcholine (PC), the major phospholipid in mammalian cells, in a concentration- and cell type-dependent manner. This generates photoswitchable PC analogs (**AzoPC**), which are predominantly located in the endoplasmic reticulum (ER). Irradiation causes a rapid photoisomerization that increases membrane fluidity with high spatiotemporal precision. We use these ‘PhotoCells’ to study the impact of membrane mechanics on protein export from the ER and demonstrate that this two-step process has distinct membrane fluidity requirements. Our approach represents an unprecedented way of manipulating membrane fluidity *in cellulo* and opens novel avenues to probe roles of fluidity in a wide variety of biological processes.

## Introduction

The ability to fine-tune membrane fluidity is essential to maintain cellular life (Ernst et al., 2016). Lipids, due to their high diversity in structure and composition (Harayama and Riezman, 2018), are the preferred tool that cells use to modulate membrane dynamics. They can function through direct protein-lipid interactions as well as indirectly through the modulation of physical membrane properties, such as bending rigidity, curvature, and fluidity. Membrane homeostasis relies on complex interdependent processes that are difficult to dissect and study in living systems with standard genetical and biochemical techniques (Dingjan and Futerman, 2021; Ernst et al., 2018). In fact, controlled manipulations of membrane fluidity have been restricted mostly to *in vitro* studies using model membrane systems, while *in cellulo*, these approaches have been limited to the use of lipid metabolic interventions that modify membrane lipid composition. Treatment of cells with polyunsaturated fatty acids (PUFAs) has been shown to facilitate membrane deformation in the context of endocytosis (Pinot et al., 2014), and increases fluidity and permeability to facilitate apoptosis (Levental et al., 2020). Similarly, modifications of lipid saturation levels have been shown to modulate mitochondrial respiration (Budin et al., 2018). These examples emphasize the importance of calibrating membrane fluidity for a proper functioning of physiological processes.

Optogenetics and photopharmacology allow for optical control of biological processes with the spatiotemporal resolution of light. While optogenetics is based on genetically encoded photoreceptors, photopharmacology relies on synthetic molecular photoswitches, such as azobenzenes (Beharry and Woolley, 2011; Hüll et al., 2018; Szymański et al., 2013). Photoswitchable lipids have emerged as versatile tools to control defined protein-membrane interactions *in vivo* (Morstein et al., 2019, 2021a) as well as membrane mechanics in model membranes(Chander et al., 2021; Doroudgar et al., 2021; Pernpeintner et al., 2017). If these lipids could be integrated into cellular membranes in sufficient quantities, they could meet a long-standing need in the field to control biophysical parameters of membranes remotely and with high spatiotemporal resolution within living systems.

Here, we engineer the lipid composition of cellular membranes using a synthetic photoswitchable fatty acid that allows us to manipulate fluidity with light. The synthetic fatty acid **FAAzo4** is efficiently incorporated into glycerophospholipids, mostly phosphatidylcholine (PC), and integrated in the endoplasmic reticulum (ER) membrane of mammalian cells. This enables us to directly modulate membrane fluidity and study the influence of this biophysical parameter on protein secretion.

## Results

### FAAzo4 is taken up by cells and integrated into AzoPC

To study the capacity of the photoswitchable lipid **FAAzo4** to be taken up by cells and metabolized to give rise to photoswitchable phospholipids (Figure 1A,B), we supplemented growth medium with analogs of **FAAzo4**, incubated HeLa cells for 4 hours, and subsequently conducted lipid extraction and mass spectrometric quantification of lipid metabolites. We envisioned that the cellular uptake of free fatty acids could be limiting for incorporation and therefore synthesized and tested a series of esterified pro-**FAAzo4** analogs. Methyl-, ethyl-, and *n*-butyl-esters are common pro-drugs for carboxylic acids (Stella et al., 2007) and acetoxymethyl esters are frequently used to mask carboxylic acids in chemical probes (Tsien, 1981). All of these pro-**FAAzo4** analogs are hydrolyzed intracellularly by non-specific esterases. While pro-**FAAzo4** (R^1^ = Me, Et, Bu, AM) analogs exhibited good cell permeability, free **FAAzo4** (R^1^ = H) yielded higher cellular levels of **AzoPC** (Figure 1C). SNAC-esters of **FAAzo4** were also prepared as metabolically activated thioester analogs of **FAAzo4** (Franke and Hertweck, 2016). They were readily taken up by cells but not incorporated into cellular phospholipids.

**Figure 1.**
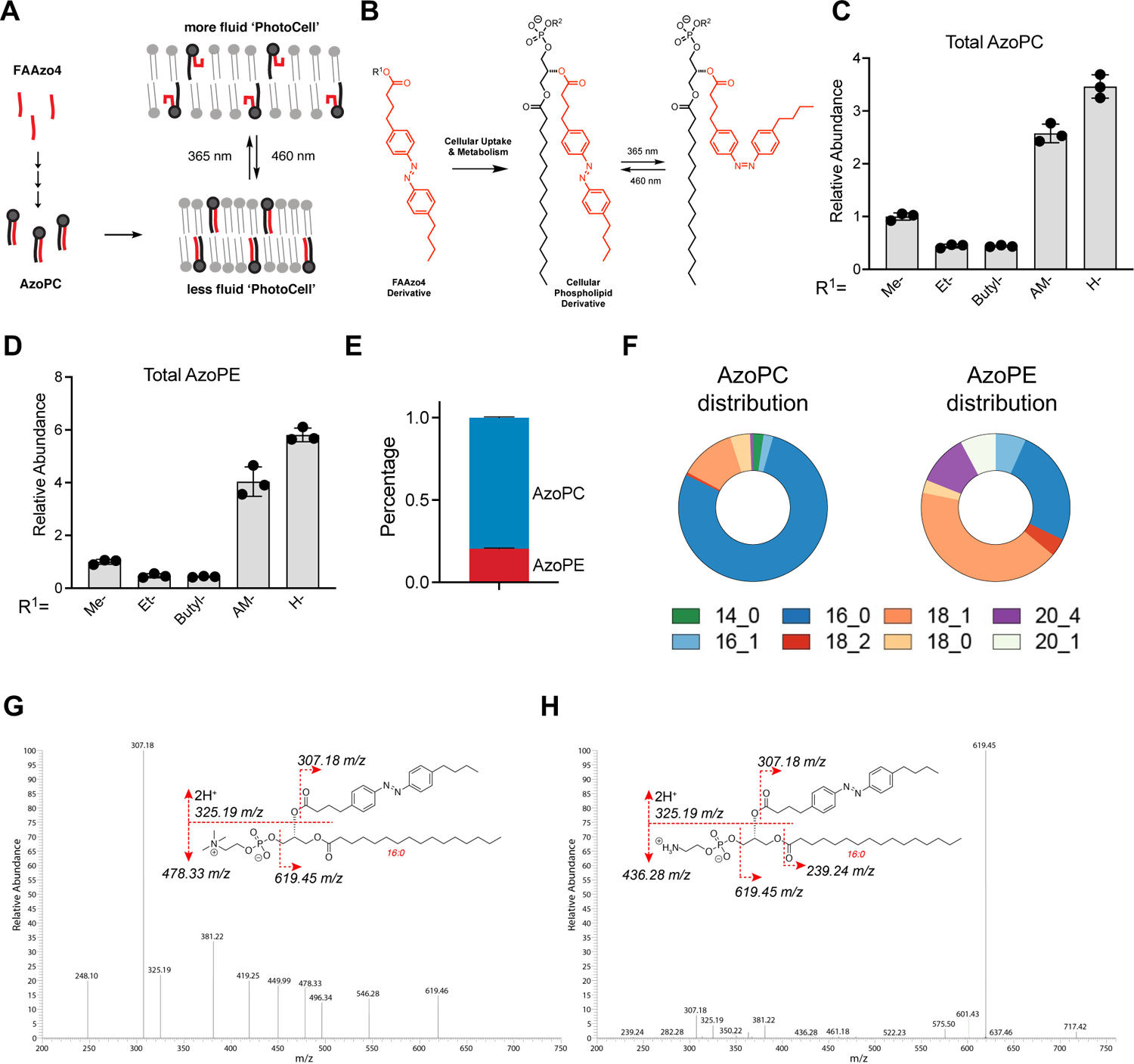
FAAzo4 is incorporated into glycerophospholipids. (A, B) Schematic illustration of PhotoCell design and photolipid structures. (C, D) Incorporation of **FAAzo4** into **AzoPC**s and **AzoPE**s after treatment of **FAAzo4** esters or **FAAzo4** in HeLa cells. (E, F) Distribution of incorporated photolipids after **FAAzo4** treatment in HeLa cells. (G,H) LC-MS/MS spectra of ions (m/z) correspond to C16:0-**AzoPC** and C16:0-**AzoPE**, respectively. Data represent three biological replicates. Error bars represent SD.

Having settled on **FAAzo4** as the best metabolic precursor, we then analyzed the composition of glycerophospholipids containing this synthetic lipid in HeLa cells. We found that **AzoPC** and photoswitchable phosphatidylethanolamine (**AzoPE**) were the predominant species (Figure 1 D,E). Interestingly, we did not detect the incorporation of **FAAzo4** into phosphatidylinositol (PI), phosphatidylserine (PS), and phosphatidylglycerol (PG), indicating selective incorporation only into the major membrane phospholipids.

Notably, incorporation into sphingolipids was also not observed. We detected *m/z* values corresponding to azobenzene-containing triacylglycerols, but their amount was negligible. To quantify the azobenzene-containing lipid metabolites, **C17-AzoPC**, comprised of heptadecanoic acid (C17:0) and **FAAzo4**, was synthesized and used as an internal standard. While **C17-AzoPC** is expected to closely mimic the ionization efficiency of **AzoPC**s, we also found that the MS profiles from **AzoPC** and endogenous PC species are similar, suggesting that the head group is the decisive factor during the ionization process in our measurement. Accordingly, the amount of **AzoPE** was determined together with endogenous PE species by the same molecule standard (PE31:1). We observed that the predominant form of **AzoPC** was C16:0, which corresponds to the chain length of the most abundant endogenous phospholipid formed. The major **AzoPE**s were C16:0 and C18:0 (Figure 1F). The presence of **AzoPC** and **AzoPE** was further confirmed by LC-MS/MS analysis (Figure 1G,H), in which the cutoff was set at *m/z* 200 to bypass the phosphocholine fragment (*m/z* 184.07), the major ion from PC species that suppresses other signals. The expected fragments of **FAAzo4** (*m/z* 307.18, 325.19) were clearly visible in all **AzoPC**s, **AzoPE**s, as well as in the reference compound **C17-AzoPC** (Figure S1A). In glycerophospholipids, the *sn-1* position is preferentially linked to saturated and monounsaturated fatty acyl chains, and the *sn-2* position is the major site for phospholipid remodeling (Wang and Tontonoz, 2019). The majority of **AzoPC**s and **AzoPE**s contains saturated and monounsaturated fatty acids (Figure 1F), suggesting that **FAAzo4** is introduced predominantly to the *sn-2* position, though we cannot rule out the possibility that the part of the acyl chain exchange occurred at *sn-1* during **FAAzo4** incorporation.

### Cellular uptake is the rate-limiting step in metabolic FAAzo4 incorporation

We next explored the mechanisms of **FAAzo4** incorporation using **AzoPC** production quantification as a readout and found that this process is dose-dependent (Figure 2A). To address if cellular uptake or metabolic incorporation are rate-limiting, we conducted a “pulse-chase” experiment, in which **FAAzo4** was washed out after 30 minutes treatment and the incorporation efficiency was measured at several time points up to four hours after the wash-out. We found that the levels of **AzoPC** did not change after the wash-out indicating that cellular uptake is the rate-limiting step for both **AzoPC** (Figure 2B) and **AzoPE** incorporation (Figure 2C), and that the incorporation likely goes through the phospholipid remodeling pathways (Wang and Tontonoz, 2019). To test if incorporation is catalyzed by members of the acyl-CoA synthetase long-chain ligase enzymes (ACSLs), we quantified levels of **AzoPC** in ACSL4-KO cells (Figure 2D) and used an ACSL inhibitor triacsin C (Tomoda et al., 1987) (Figure 2E). In both cases we found that the levels of **AzoPC** formed in PhotoCells was reduced compared to the control, indicating the involvement of ACSLs during the incorporation. We also observed that *trans*-**FAAzo4** was incorporated more efficiently than *cis*-**FAAzo4** (Figure 2F), which was obtained through pulsed irradiation with 370 nm light (75 ms every 15 s) using a Cell DISCO system (Borowiak et al., 2015; Morstein and Trauner, 2020). The amount of **AzoPC** could be markedly increased through serum starvation depleting medium of other fatty acids (Figure 2G). The viability of PhotoCells was tested under our standard incorporation conditions (Figure S1B). We found that the cell viability was not compromised, indicating that **AzoPC** was well tolerated in membranes of live cells.

**Figure 2.**
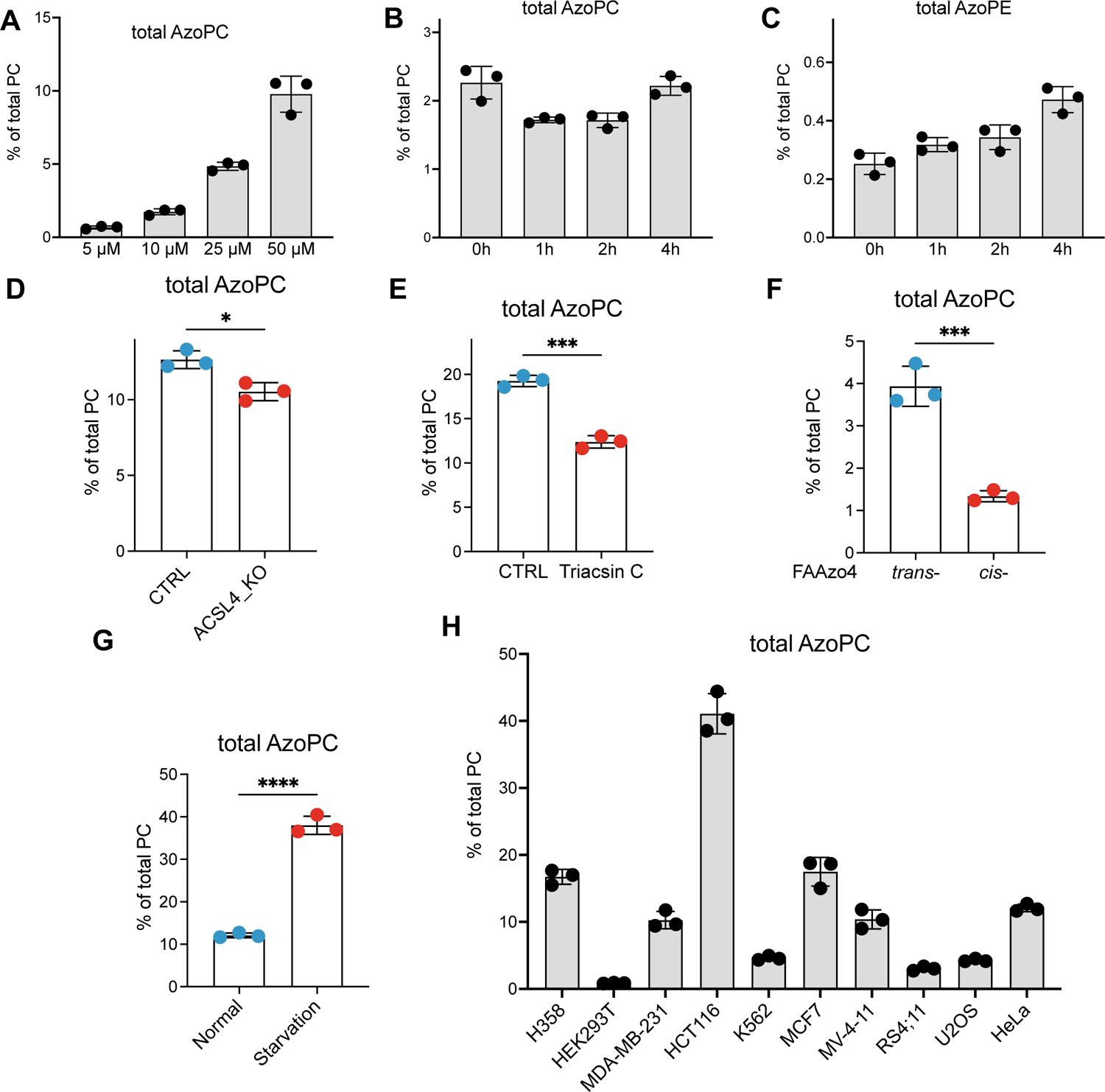
Quantification of FAAzo4 incorporation under various experimental conditions. Cells were treated with 50 μM **FAAzo4** for 4 hours unless otherwise stated. Levels of **AzoPC** were quantified to measure **FAAzo4** incorporation efficiency. (A) Concentration dependence of incorporation. HeLa cells were incubated with **FAAzo4** at indicated concentrations. (B, C) HeLa cells were incubated with **FAAzo4** for 30 minutes, replaced by fresh medium, and then collected at indicated time points. (D) HeLa cells were incubated with **FAAzo4** in control or ACSL4 KO cells. (E) HeLa cells were incubated with **FAAzo4** in the presence/absence of 10 μM Triacsin C. (F) HeLa cells were incubated with *trans-* or *cis-* **FAAzo4**. (G) After 24-hour starvation, HeLa cells were incubated with **FAAzo4** and compared to cells in normal growth conditions. (H) Incorporation efficiency of **FAAzo4** in different cell lines. Data represent three biological replicates. Error bars represent SD, *p<0.05, **p<0.01, ***p<0.001, students’ *t*-test.

To test the generality of our PhotoCell approach, we tested **FAAzo4** incorporation in a range of adherent and suspension cell lines (Figure 2H). Several cell lines showed effective incorporation yielding 10-20% **AzoPC**, including H358, MDA-MB-231, MCF7, MV-4-11, and HeLa cells. Interestingly, HEK293T cells do not exhibit effective incorporation, whereas HCT116 cells showed exceptional incorporation yielding up to 40% **AzoPC**. Based on these results, we decided to use HeLa and HCT116 PhotoCells in subsequent experiments.

### Remodeled AzoPC is Predominantly Located in ER Membranes

We next examined the subcellular localization of *in cellulo* formed photolipids in HCT116 cells to determine which membranes could be studied with this approach. Since ER membranes are known to be the major site of phospholipid remodeling, (Wang and Tontonoz, 2019) we hypothesized that **FAAzo4** is predominantly incorporated into glycerophospholipids at this site. To test if the localization of **AzoPC** is confined to ER membranes, we employed a clickable analog of **FAAzo4**, termed **clFAAzo4** (Figure 3A). Through minimal chemical modification with a terminal alkyne, Copper-Catalyzed Azide-Alkyne Cycloaddition (CuAAC) can be used in fixed cells to visualize the location of photolipid metabolites (Figure 3B). The soluble fluorophore Sulfo Cy5 is ideally suited for this experiment as its high water solubility prevents accumulation in membranes, which can be overcome through conjugation to a lipid with two acyl tails (Morstein et al., 2021b; Walter et al., 2017). Subsequent addition of an ER marker enabled co-localization. This experiment showed strong overlap of **AzoPC** with membranes of the ER (Pearson Coefficient 0.95, Figure 3C), suggesting that PhotoCells could be particularly suited to the study of ER membrane biophysics. The ER membrane naturally exhibits high levels of phosphatidylcholine and lower levels of the major plasma membrane lipids cholesterol and sphingomyelin (van Meer et al., 2008).

**Figure 3.**
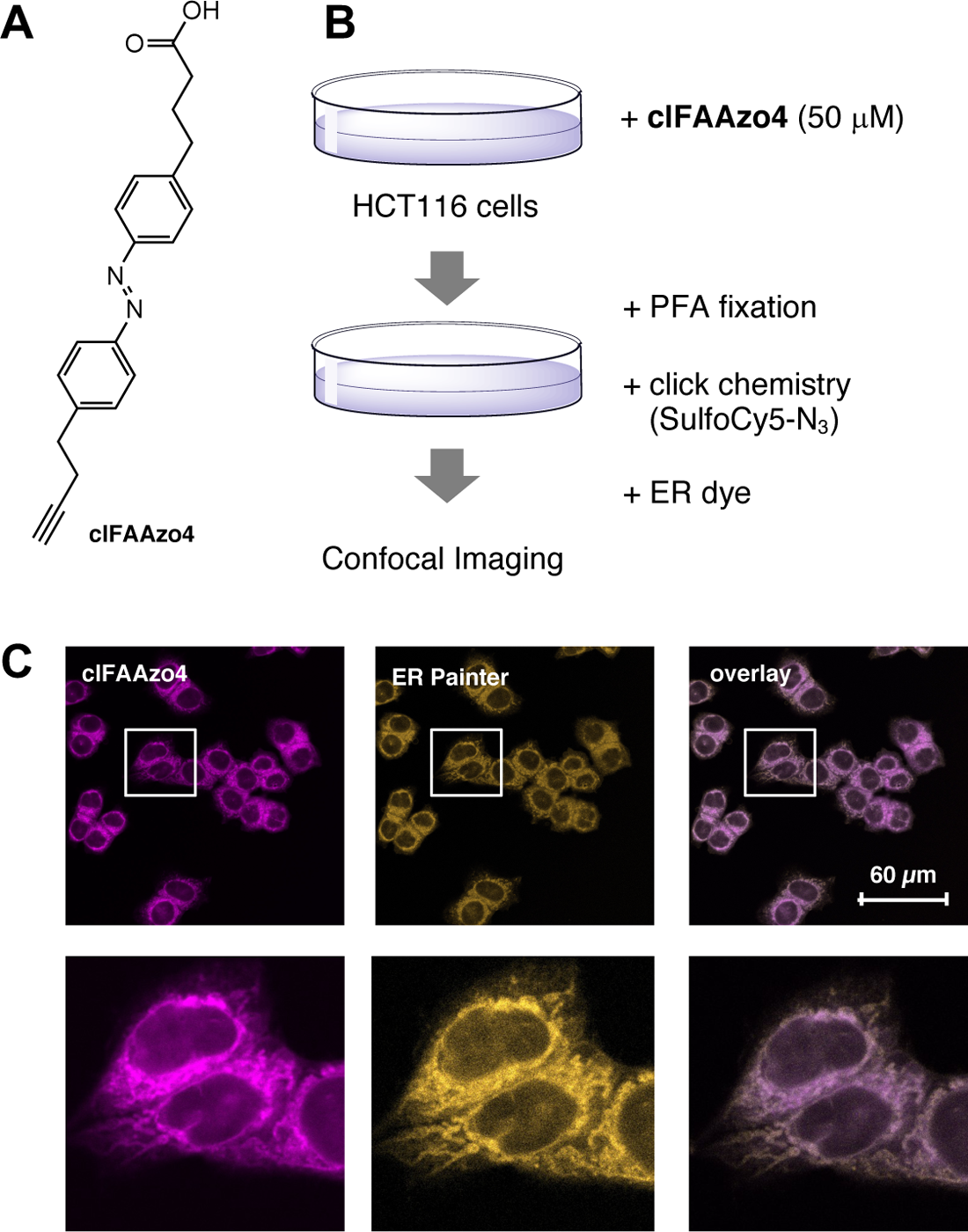
Incorporation and click-imaging of alkyne-modified FAAzo4. (A) Chemical structure of clickable analog **clFAAzo4**. (B) Schematic of click-imaging protocol for incorporated **clFAAzo4** in fixed cells. HTC116 cells were incubated with **clFAAzo4**, fixed with para-formaldehyde, and then labeled with sulfo-cyanin-5-azide by means of CuAAC. Subsequently, cells were treated with a fixation-compatible ER-selective dye. (C) Confocal images (63 x) were obtained using *λ*_Ex_ = 646 nm for Cy5 and *λ*_Ex_ = 488 nm for ER Painter.

### C16-AzoPC Enables Optical Control of Membrane Fluidity *in Vitro*

To study the effect of the predominant metabolite of **FAAzo4**, **C16-AzoPC,** on membranes in more detail, we chemically synthesized **C16-AzoPC** and performed measurements on two common model systems, namely supported lipid bilayers (SLBs) and giant unilamellar vesicles (GUVs) (Figure 4). We previously reported that photoisomerization of photolipid model membranes containing **C18-AzoPC** impacts the membrane fluidity (Urban et al., 2020) (Figure 4A). We observed similar effects with **C16-AzoPC**. The fluidity was investigated by performing fluorescence recovery after photobleaching (FRAP) measurements on SLBs containing 99 mol% **C16-AzoPC** and 1 mol% of dye-labeled lipids (TexasRed-DHPE). Imaging and photoswitching of the **C16-AzoPC** SLB were simultaneously achieved by using a UV-A (330-385nm) and green (510-550nm) filter cube. The lateral diffusion was then determined by localized bleaching of a small bilayer area and the subsequent analysis of the fluorescence recovery (Figure 4B). The average diffusion coefficient of a *trans*-adapted SLB is D*_trans_* = (0.37±0.02) µm^2^s^-1^ (Figure 4C, left). After changing the illumination to UV-A light, the diffusion increases by a factor of ∼4. The average lateral diffusion coefficient of a *cis*-adapted SLB is D*_cis_* = (1.62±0.06) µm^2^s^-1^, which is indicative for a fluid bilayer membrane (Almeida and Vaz, 1995). This increase in fluidity can be explained by a decrease of lipid-lipid interactions between photolipids in the *trans* state that can hinder lateral lipid diffusion, as we showed previously for **C18-AzoPC** membranes (Urban et al., 2018, 2020). We further performed temperature dependent FRAP experiments and heated a **C16-Azo-PC** SLB in steps of ∼10°C up to 60°C (Figure 4C, right). The lateral diffusion coefficients of the *cis*-**C16-AzoPC** membrane remained constant over the investigated temperature range of 21-60°C. The fluidity of the *trans*-**C16-AzoPC** SLB increases from ∼0.4 µm^2^s^-1^ to ∼0.7 µm^2^s^-1^, indicating an H-aggregate melting. In a next step, we investigated if these effects are also retained in more native model membranes. We therefore added the abundant endogenous PC analog POPC and repeated the diffusion experiments with SLBs consisting of a 1:1 mixture of these lipids. This ratio corresponds to the absolute amount of **C16-AzoPC** obtained in HCT116 or starved HeLa MZ PhotoCells. The average lateral diffusion coefficient increases from D*_trans_*= (0.93±0.05) µm^2^s^-1^ to D*_cis_*= (1.28±0.05) µm^2^s^-1^ upon *trans*-to-*cis* isomerization (Figure 4D, left). The magnitude is less compared to the **C16-AzoPC** SLB, which can be explained by a reduction of the dipolar interactions upon dilution of the **AzoPC**-aggregates by the regular non-switchable lipid POPC. A comparable effect was found for membrane mixtures of **C18-AzoPC** and DPhPC (Urban et al., 2018). This is also supported by the small increase of the lateral diffusion from ∼0.8 µm^2^s^-1^ to ∼0.9 µm^2^s^-1^ upon heating the sample to 60°C (Figure 4D, right). The change of the membrane fluidity upon photoswitching further suggests a strong influence on the mechanical properties of the **C16-AzoPC** containing membranes. We therefore performed fluorescence microscopy of cell-sized GUVs that were labeled with 1mol% TexasRed-DHPE (Figure 4E&F). In the *trans* state, the GUVs show a spherical shape. On exposure with UV-A light, strong membrane fluctuations arise within only a few milliseconds, which stop again immediately upon green light illumination. This observation implies that *trans*-to-*cis* photoswitching results in a decrease of the vesicle stiffness, which is in line with a fluidity increase as derived from the FRAP data and also with pervious results of **C18-AzoPC** GUVs (Pernpeintner et al., 2017). Altogether, the findings confirm that the observed photoswitching effects on the photolipid membrane properties are consistent at different concentrations of **C16-AzoPC** and dose-dependent.

**Figure 4.**
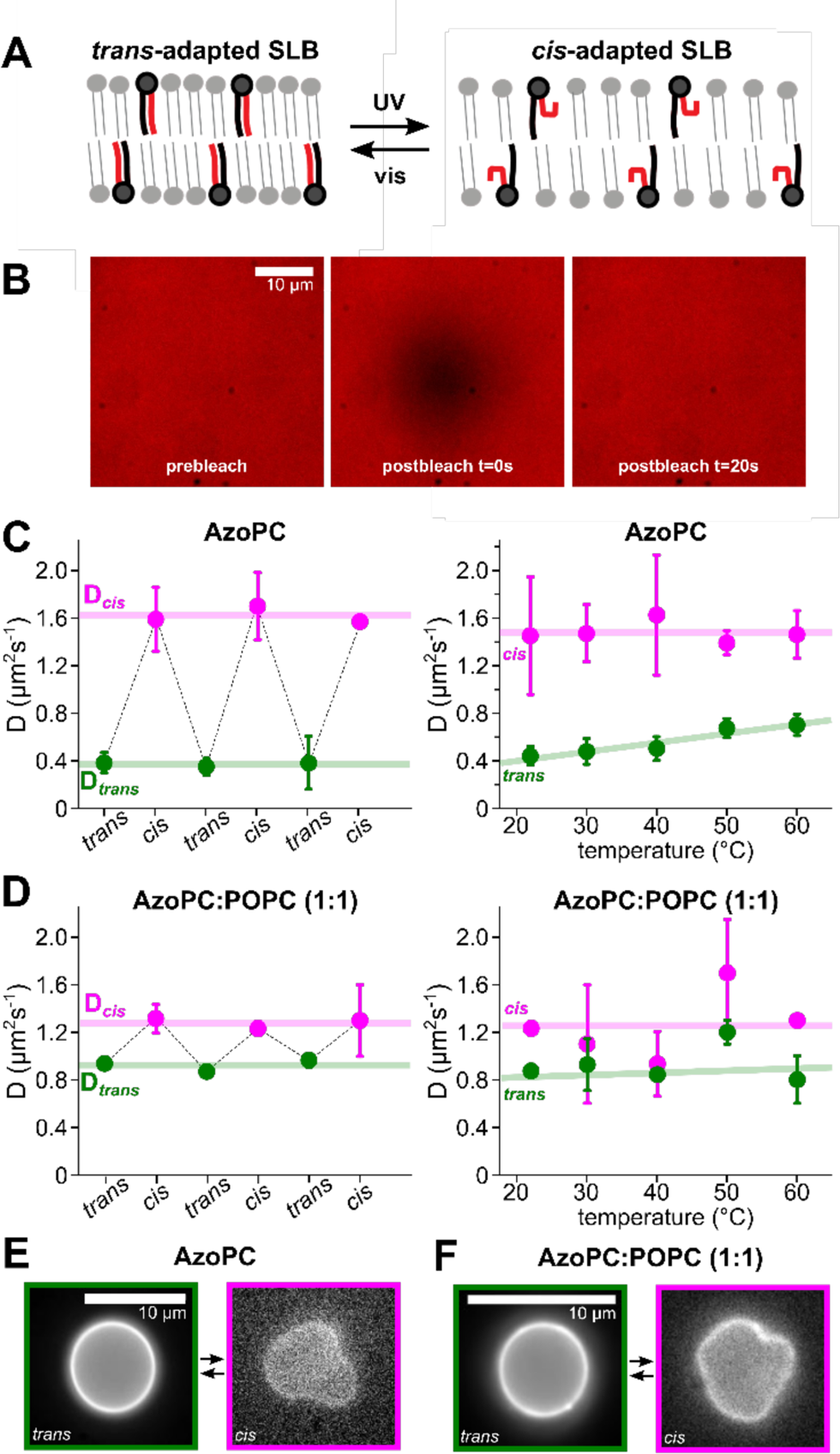
Biophysical modulation of C16-AzoPC containing model membranes. (A) Schematic depiction of light-induced membrane fluidity modulation with **AzoPC**. (B) Fluorescence images of a SLB, illustrating the bleach spot and fluorescence recovery of a FRAP experiment. (C) Diffusion coefficients of the **C16-AzoPC** SLB. (D) Diffusion coefficients of the **C16-AzoPC**:POPC (1:1) SLB. (E,F) Fluorescence images of dye labeled GUVs. Switching between *trans* and *cis* changes the vesicle shape and membrane fluctuations.

### ER Membrane Fluidity Controls Cargo Export

To provide further evidence that the plasma membrane is not affected, we explored the effects of photolipids on signal transduction cascades and viral infection. Phosphoproteomics was used to read out cellular signaling processes mediated by GPCRs, RTKs, and other membrane receptors. No meaningful change in the phosphoproteome was detected upon PhotoCell irradiation (Figure S2). We also did not observe a light-dependent effect on vesicular stomatitis virus (VSV) infection, a commonly used model to examine cell entry via endocytosis (Figure S3). This suggests that our system can be specifically used for the study of ER dependent processes.

One of the major functions of this organelle is the process of protein export by the COPII machinery (Brandizzi and Barlowe, 2013). This process involves sequential protein recruitments to the ER membrane followed by membrane bending and fission to give rise to COPII carriers that transport proteins from the ER to the Golgi. This operation has been previously shown to be modulated by lipid composition, which affects both lipid-protein interactions and membrane mechanical properties necessary for bending and fission (Contreras et al., 2012; Jiménez-Rojo et al., 2020; Melero et al., 2018; Rodriguez-Gallardo et al., 2020). In order to study the contribution of membrane fluidity on this process we took advantage of the Retention Using Selective Hooks (RUSH) system, an approach to synchronize and follow the export of proteins from the ER (Boncompain et al., 2012). We used two different constructs, mCherry-TNF*α* (TNF*α*-RUSH) and EGFP-GPI (GPI-RUSH) and followed the secretion of these proteins using high-content microscopy (Fig. 5 A, B).

**Figure 5.**
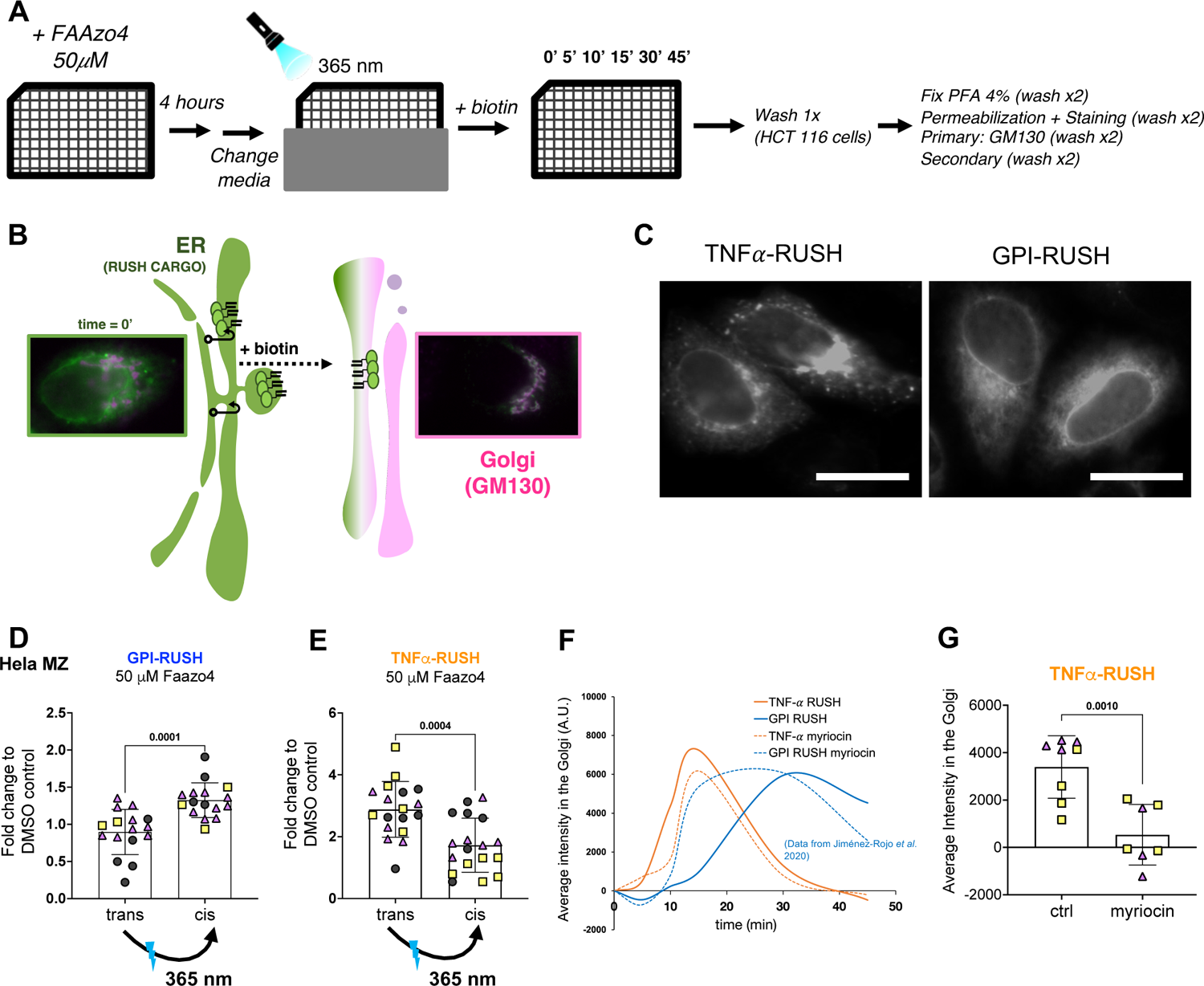
Optical control of membrane mechanics followed by RUSH reveals sequential fluidity requirements for protein secretion. (A) Scheme describing the experimental set up. **FAAzo4** treatment and “Retention using selective hooks” assay was performed in 96-well plates to allow High Content microscopy. After **FAAzo4** treatment cells were illuminated with UV-A light at 365 nm and control wells were covered with aluminum foil. Biotin was added at different time points to release the transfected cargos and after the experiment the cells were fixed and prepared for staining and imaging. (B) Schematic representation of the RUSH assay. (C) Representative images showing expressed selected cargo in HeLa cells before release upon biotin addition. (D) Effect of light-induced increase in membrane fluidity on the secretion of the GPI-RUSH construct in HeLa cells. Each point represents the average of around 500 individual cells. Time = 15 minutes after biotin addition. (E) Effect of light-induced increase in membrane fluidity on the secretion of the TNF*α*-RUSH construct in HeLa cells. Time = 10 minutes after biotin addition. (F) Representative curves showing secretion dynamics of GPI-RUSH construct and TNF*α*-RUSH in control cells and in cells treated with the inhibitor of sphingolipid synthesis myriocin. Data for GPI-RUSH construct was taken from (Jiménez-Rojo et al., 2020). (G) Effect of myriocin treatment on the secretion of TNF*α*-RUSH construct in HeLa cells. Statistical analysis was performed by unpaired two-tailed Student’s t test with Welch’s correction.

Thus, combining our approach, which allows for optical control of membrane fluidity, with the RUSH assay, we are able to evaluate the contribution of membrane fluidity to the secretion of TNF*α* and a model GPI-anchored protein (Fig 5A,C). Notably, the secretion of GPI-RUSH was accelerated after photoswitching cellular **AzoPC** to *cis* leading to increased membrane fluidity. Opposing effects were observed for TNF*α*-RUSH, where export is decelerated after irradiation and the subsequent increase of membrane fluidity (Fig 5D, E).

In a separate experiment, we increased membrane fluidity using the sphingolipid synthesis inhibitor myriocin (Fig S4D), which has previously been shown to accelerate the transport of GPI-RUSH (Fig 5F) (Jiménez-Rojo et al., 2020). In agreement with our findings in PhotoCells, the transport of TNF*α*-RUSH is halted by myriocin treatment, thus independently confirming the distinct membrane fluidity requirements for the secretion of the two cargos. Notably, before biotin addition (time=0) the GPI-RUSH protein is homogeneously dispersed through the ER, while TNF*α*-RUSH is located in discrete regions that probably correspond to ER exit sites (ERES), as previously described by Weigel *et al*. (Weigel et al., 2021) (Fig. 5C, S4C). This is supported by the fact that the export of TNF*α*-RUSH to the Golgi started already 5 minutes after release (Fig. 5F, S4B). As such, the export of the two different cargos might be representative of two different steps of the protein secretion process: first, accumulation of the protein et the ERES and second, export to the Golgi. Taken together, our data support distinct membrane fluidity requirements for the secretion of two different proteins from the ER that could reflect differential fluidity requirements for these two steps in protein export.

### The biological effects of optical membrane modulation depend on the amount of photolipids

To evaluate the concentration dependency of **AzoPC**-induced biological effects, in particular the impact on protein export kinetics, we applied our RUSH assay in HCT 116 cells where the incorporation of **FAAzo4** into PC phospholipids at 50 μM was around 40% (Figure 2H). To assess the incorporation efficiency of **FAAzo4** in HCT116 cell line, we treated the cells with different concentrations of **FAAzo4** and found that treatment with 10 μM of **FAAzo4** in HCT 116 would yield levels of **AzoPC** equivalent to treatment with 50 μM **FAAzo4** in HeLa cells (Fig 2H, Fig 6A). In agreement with this, we found that after treating HCT 116 cells with 10 μM **FAAzo4**, the effect on transport kinetics of GPI-RUSH went in the same direction as what we demonstrated for HeLa cells. However, as we push the incorporation of **FAAzo4** to higher levels, the effect seems to be detrimental rendering transport kinetics of GPI-RUSH slower. To test if higher levels of **AzoPC** lead to ER stress, we analyzed RNA levels of CHOP as a read-out of the activation of unfolded protein response (UPR). Indeed, we found that levels of 25% **AzoPC** and higher led to ER stress and UPR activation, which hinders the interpretation of our transport assays (Fig 6C). We thus conclude that, in order to use **FAAzo4** for cell biology assays, the amount of its incorporation should be carefully monitored and optimized in a given biological system. Cells that have 10% of their cellular PC substituted with **AzoPC**, (i.e., HeLa cells treated with 50 μM **FAAzo4** or HCT 116 cells treated with 10 μM **FAAzo4**) appear to be physiologically unaltered and offer a more relevant background for the study of cellular processes, while cells that have 40% of their PC in the form of **AzoPC** undergo ER stress (Fig 6C, D). This fits well with our observation that 10 μM treatment in HCT 116 cells has little effect on the overall lipidome of the cells, while 50 μM leads to the formation of several secondary metabolites of cellular phospholipids (Fig S5A-C).

**Figure 6.**
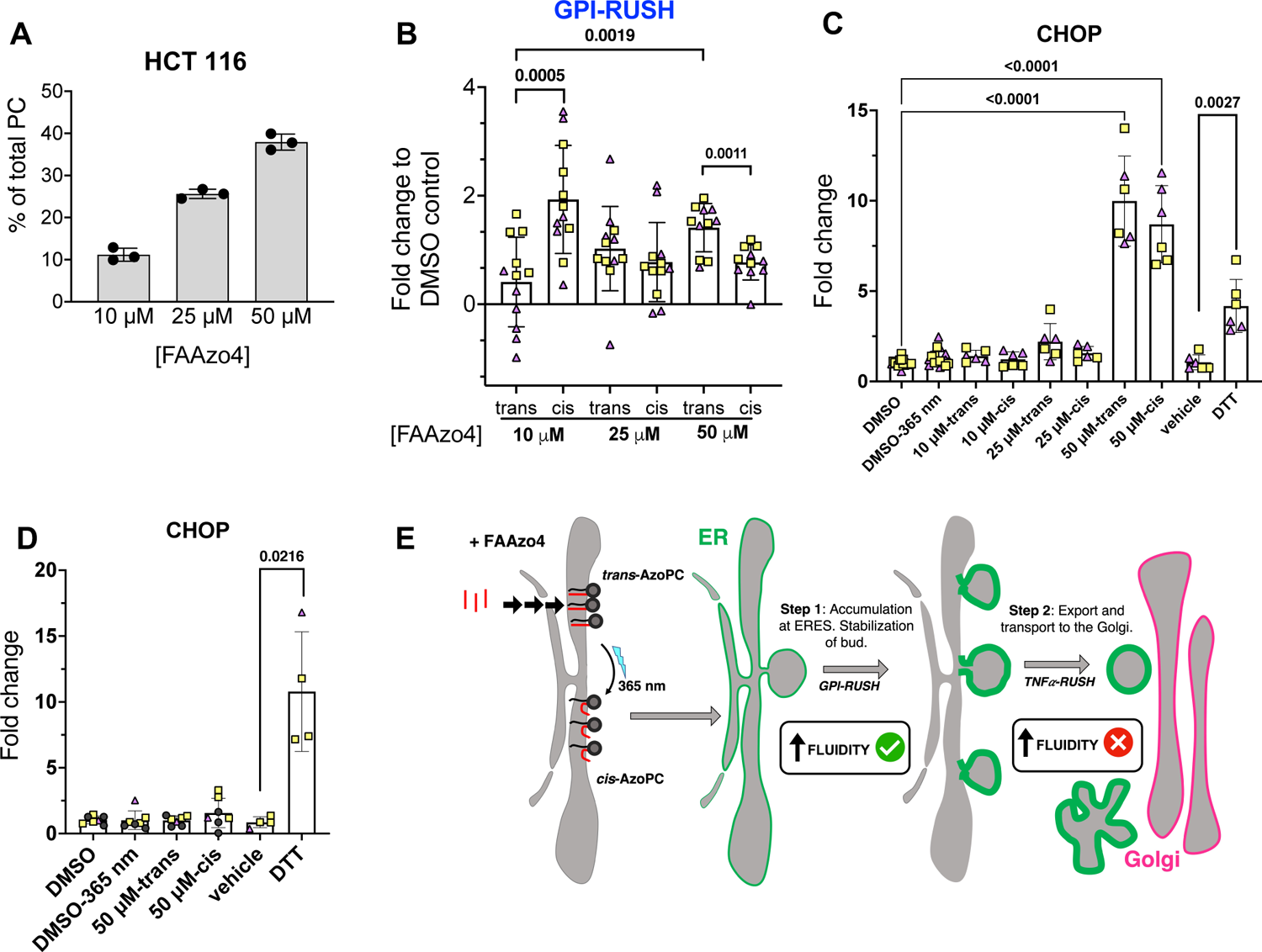
The physiological effects of photolipids are concentration dependent. (A) Incorporation of **FAAzo4** in HCT 116 cells as measured by mass spectrometry. (B) Effect of light-induced increase in membrane fluidity on the secretion of the GPI-RUSH construct in HCT 116 cells treated with different amounts of **FAAzo4**. (C) Expression of the ER stress marker CHOP upon **FAAzo4** treatment in HCT 116 cells before and after irradiation. (D) Expression of the ER stress marker CHOP upon 50 μM **FAAzo4** treatment in HeLa cells before and after irradiation. (E) Working model.

## Discussion

In recent years, photoswitchable lipids have been used increasingly for the control of lipid signaling pathways and membrane properties (Morstein et al., 2021a). While the former was often done in live cells, the modulation of membrane mechanics with photolipids was exclusively studied in reconstituted systems (e. g. GUVs and SLBs). We now show that photolipids can be integrated into membranes of live cells in a surprisingly simple fashion. This offers unprecedent opportunities to study the contribution of membrane properties, such as its fluidity, to cell physiology. Photoswitchable lipids have the advantage that they preserve the integrity of the headgroups of endogenous lipids, allowing them to function similarly to their native analogs, and they can rapidly change the physicochemical properties of their lipophilic part.

The engineering of PhotoCells was achieved through the feeding of a simple photoswitchable fatty acid analog **FAAzo4**. This precursor is effectively metabolized into phospholipids, mostly **AzoPC**, which is predominantly localized at the endoplasmic reticulum in a variety of cell lines.

Having established the effective and selective incorporation of **AzoPC** in the ER membrane, we demonstrated that our approach can be readily combined with other recently developed cell biology techniques to study membrane trafficking. Membrane fluidity and curvature have been proposed to affect several ER-localized processes, such as protein translocation and lipid droplet assembly (Nilsson et al., 2001; Santinho et al., 2020). Moreover, protein secretion by COPII is known to be modulated by lipid-protein interactions and very likely by changes in membrane curvature, tension, and asymmetry.

In this context, lysolipids have been proposed to facilitate COPII vesicle formation by decreasing the energy required for membrane bending (Melero et al., 2018). Sphingolipids and ether lipids have been shown to play a role in the secretion of GPI-anchored proteins through specific lipid-protein interactions and likely through affecting membrane biophysical properties (Jiménez-Rojo et al., 2020; Rodriguez-Gallardo et al., 2020). Additionally, the protein export mechanism differs depending on the selected cargo. It is known that the export of bulky cargos, such as GPI anchored proteins, requires specific COPII protein isoforms and it is also regulated by the cargo crowding itself (D’Arcangelo et al., 2015; Gomez-Navarro et al., 2020; Muñiz et al., 2000). Cargos like pro-collagen need distinct adaptors that support their transport into very large non-canonical carriers (TANGO1, cTAGE5) and modulate membrane tension to facilitate elongation of the bud and of transport by acquiring a ring-like structure (Raote et al., 2018, 2020). Furthermore, a two-step secretion model has recently been proposed by Weigel *et al*. where they show that the accumulation of the cargo at ERES and export to the Golgi are two independent processes. Moreover, they were able to measure the kinetics of both steps using different cargos (Weigel et al., 2021). First, they showed, using the RUSH technology, that TNF*α*-RUSH is already located at ER exit sites (ERES) before the release of the cargo and that the first minutes after biotin addition report on the export of the protein to the Golgi. By contrast, in the case of GPI-RUSH, the first minutes after release would report on the accumulation of the cargo at ERES. The approach we describe here allowed us to rapidly modify membrane fluidity in the ER and to assess the respective membrane fluidity requirements. Increased membrane fluidity accelerated cargo accumulation at ERES, presumably through facilitating protein diffusion. In addition, we showed that increased membrane fluidity decreased the rate of ER cargo export. Due to the high temporal resolution of this approach, we were thus able to establish membrane fluidity as direct contributor to protein export. Our data support the two-step secretion model and provide insights into the role of membrane fluidity at each step.

The ER is the home of many other cellular processes, including protein folding, protein quality control, the unfolded protein response UPR), ER associated degradation (ERAD), lipid biosynthesis, lipid droplet formation, and the ER forms contact sites with all of the other organelles in the cell. PhotoCells should be applicable to the study of membrane fluidity in these processes and, potentially, in other organelles (see limitation section). They represent a novel approach to gaining direct insight into the role of membranes in biological processes using light as a non-invasive input signal that affords high spatiotemporal control. As such, our approach complements optogenetic methods, which are based on the expression of photoreceptor proteins, and have already been widely employed in cell biology (Deisseroth, 2011; Fenno et al., 2011).

### Limitations of this Study

The use of UV-A light for photoswitching (365 nm) restricts the utility of our approach *in vivo* where penetration of light with this wavelength could be a limiting factor. Future studies will therefore involve red-shifted photolipids to be able to employ red or near-IR light with higher tissue penetration.

Secondly, the described assay is limited to functional studies in the ER due to preferential localization of **AzoPC** to the ER. Future studies should address methods to target **AzoPC** to other organelles. This could be based on differential photolipids precursors (e.g. using organelle-targeting groups) or heterologous expression of enzymes that control phospholipid metabolism in other organelles.

Furthermore, the usefulness of PhotoCells depends on the level of metabolic incorporation. In HCT 116 cells, where the incorporation is more efficient, the high amounts of **AzoPC** can induce ER stress and activate UPR providing a non-reliable background to study cellular processes. Therefore, it is important to monitor and optimize the levels of **AzoPC** depending on the biological system used.

## Materials and Methods

### EXPERIMENTAL MODEL AND SUBJECT DETAILS

#### Cell lines

The Human Acute Lymphoblastic leukemia RS4;11 (ATCC CRL1873) cell line and the human biphenotypic B-myelomonocytic leukemia MV4-11 (ATCC CRL-9591) cell line were purchased from the American Type Culture Collection (ATCC) and cultured in RPMI 1640 medium ( Cat# 11835030, Gibco, no phenol red) supplemented with 10% fetal bovine serum and 1% penicillin/streptomycin (Cat# 15140122) in a humidified incubator at 37°C with 5% CO_2_ in air.

Hela MZ cells were a gift from Prof. Jean Gruenberg (University of Geneva) and were cultured at 37°C and 5 % CO_2_ in Dulbecco’s Modified Eagle Medium (GLUTAMAX, DMEM, GIBCO^TM^) with 4.5 g/L glucose, supplemented with 10 % fetal calf serum (FCS, Cat# 10270106, Gibco), or in FluoroBrite DMEM supplemented with 10 % FCS, 1 % Pen/Strep when UVA light is applied to the cells.

HCT 116 cells were cultured at 37°C and 5 % CO_2_ in FluoroBrite DMEM (Cat# A1896701, DMEM, Gibco) with 4.5 g/L glucose, supplemented with 10 % fetal calf serum (FCS, Cat# 10270106, Gibco), 1 % Pen/Strep (Cat#10378016) and Glutamine (Cat# 25030024, GIBCO^TM^).

MDA-MB-231 were a gift from Prof. Michele Pagano (New York University, originally obtained from ATCC) were cultured at 37 °C and 5% CO_2_ in DMEM (high glucose 4.5 g/l; Gibco, Thermo Fisher cat# 10566024) supplemented with 10% FBS (Gibco, Thermo Fischer cat# 10437036), 1% penicillin-streptomycin-glutamine (Gibco, Thermo Fischer cat# 10378016) and a final concentration of 4 mM L-glutamine (Gibco, Thermo Fischer cat# 25030081).

Human embryonic kidney 293T (HEK 293T) cells (ATCC CRL-3216) were cultured at 37 °C and 5% CO_2_ in DMEM (Gibco) supplemented with 4.5 g/L glucose, 10% FBS (Gibco), 1% penicillin-streptomycin-glutamine and a final concentration of 4 mM L-glutamine (Gibco).

U2OS osteosarcoma cells and HCT 116 colorectal carcinoma cells were a gift from Prof. Michele Pagano (New York University, originally obtained from the ATCC) and were cultured at 37 °C and 5% CO_2_ in McCoy’s 5A medium (Cat# 16600082, Gibco) supplemented with 10 % fetal calf serum (FCS, Cat# 10270106, Gibco) and 2 mM L-glutamine, 100 IU/mL penicillin, and 100 μg/mL streptomycin (Cat# 10378016, Gibco).

K-562 Chronic Myelogenous Leukemia cells (ATCC, CCL-243) were cultured at 37 °C and 5% CO_2_ in IMDM (Cat# 12440053, Gibco) supplemented with 10 % fetal calf serum (FCS, Cat# 10270106, Gibco) and 2 mM L-glutamine, 100 IU/mL penicillin, and 100 μg/mL streptomycin (Cat# 10378016, Gibco).

MCF7 cells were a gift from Prof. Michele Pagano (New York University, originally obtained from the ATCC) and were cultured at 37 °C and 5% CO_2_ in Dulbecco’s Modified Eagle Medium (DMEM, Gibco) supplemented with 10 % fetal calf serum (FCS, Cat# 10270106, Gibco) and 2 mM L-glutamine, 100 IU/mL penicillin, and 100 μg/mL streptomycin (Cat# 10378016, Gibco).

## METHOD DETAILS

### General Methods

All reagents and solvents were purchased from commercial sources (Sigma-Aldrich, TCI Europe N.V., Strem Chemicals, etc.) and were used without further purification unless otherwise noted. Reactions were monitored by TLC on pre-coated, Merck Silica gel 60 F_254_ glass backed plates. Flash silica gel chromatography was performed using silica gel (SiO_2_, particle size 40-63 μm) purchased from Merck. All NMR spectra were measured on a BRUKER Avance III HD 400 (equipped with a CryoProbe^TM^). Multiplicities in the following experimental procedures are abbreviated as follows: s = singlet, d = doublet, t = triplet, q = quartet, quint = quintet, sext = sextet, hept = heptet, br = broad, m = multiplet. Proton chemical shifts are expressed in parts per million (ppm, *δ* scale) and are referenced to the residual protium in the NMR solvent (CDCl_3_: *δ* = 7.26 CD_3_OD: *δ* = 3.31). Carbon chemical shifts are expressed in ppm (*δ* scale) and are referenced to the carbon resonance of the NMR solvent (CDCl_3_: *δ* = 77.16; CD_3_OD: *δ* = 49.00). High-resolution mass spectra (HRMS) were obtained with an Agilent 6224 Accurate Mass time-of-flight (TOF) LC/MS system using either an electrospray ionization (ESI) or atmospheric pressure chemical ionization (APCI) ion source. All reported data refers to positive ionization mode.

### General Lipid Feeding Protocol

Cells were maintained in T25 flasks with medium containing 10 % fetal bovine serum and 1 % penicillin/streptomycin at 37 °C in a humidified 5 % CO_2_ atmosphere. After the cells were seeded and incubated overnight, the medium was carefully removed with a pipette without disturbing the cells. Cells were incubated with **FAAzo4** (10-50 µM in growth medium, prepared from 50 mM DMSO stock) at 37 °C and 5 % CO_2_ for 4 hours or at indicated time.

### Cell Viability Experiment

To assess cell viability, Hela cells were seeded in a 96 well plate (approx. 10k per well in 100 µL) in DMEM/FCS/PS (90:10:1). After 24 h cells were fed with **FAAzo4** according to the standard cell feeding protocol for 4 h and DMSO was used as a control. After 24 h of compound incubation PrestoBlue (Thermo Scientific) was added (20 µL per well) and after 2h fluorescent was measured using a BMG Labtech FLUOstar Omega microplate reader with 544/590 nm filters.

### Lipid Extraction

Lipids were extracted following previously described protocols.(Guan et al., 2009) Briefly, cells were washed by cold PBS and scraped off in 500 μL cold PBS on ice. The suspension was transferred to a 2.0 mL Eppendorf Safe-Lock Tube in which it was spin down at 2,500 rpm for 5 minutes at 4 °C. After carefully taking off the PBS, samples were extracted following the MTBE protocol(Matyash et al., 2008). Briefly, cells were re-suspended in 100 μL of water, 360 μL of MeOH and a mixture of internal standards (1 nmol of C17AzoPC, 0.4 nmol of DLPC, 1 nmol of PE31:1, 1 nmol of PI31:1, 3.3 nmol of PS31:1, 2.5 nmol of C12 sphingomyelin, 0.5 nmol of C17 ceramide and 0.1 nmol of C8 glucosylceramide) was added. Samples were vortexed, following the addition of 1.2 ml of MTBE. The samples were vigorously vortexed at maximum speed for 10 minutes at 4 °C and incubated for 1 h at room temperature on a shaker. Phase separation was induced by addition of 200 μL MS-grade water and incubation for 10 minutes. Samples were centrifuged at 1,000 g for 10 minutes. The upper phase was transferred into a 13 mm glass tube and the lower phase was re-extracted with 400 μL of a MTBE/MeOH/H2O mixture (10:3:1.5, v/v). The extraction was repeated one more time. The combined organic phase was dried by nitrogen flow.

For analysis of the full lipidome, the MTBE extract was divided, and one aliquot was deacylated to eliminate phospholipids by methylamine treatment (Clarke method). 0.5 mL monomethylamine reagent (MeOH/H2O/n-butanol/Methylamine solution (4:3:1:5 v/v) was added to the dried lipid, followed by sonication (5 min). Samples were then mixed and incubated for one hour at 53°C and dried (as above). The monomethylamine treated lipids were desalted by n-butanol extraction. 300 μl H_2_O saturated n-butanol was added to the dried lipids. The sample was vortexed, sonicated for 5 min and 150 μl MS grade water was added. The mixture was vortexed thoroughly and centrifuged at 3200 x g for 10 min. The upper phase was transferred in a 2 mL amber vial. The lower phase was extracted twice more with 300 μl H_2_O saturated n-butanol and the upper phases were combined and dried. TL and SL aliquots were resuspended in 250 μl Chloroform/methanol (1:1 v/v) (LC-MS/HPLC GRADE) and sonicated for 5 minutes. The samples were pipetted in a 96 well plate (final volume = 100 μl). The TL were diluted 1:4 in negative mode solvent (Chloroform/Methanol (1:2) + 5mM Ammonium acetate) and 1:10 in positive mode solvent (Chloroform/Methanol/Water (2:7:1 v/v) + 5mM Ammonium Acetate). The SL were diluted 1:10 in positive mode solvent and infused onto the mass spectrometer. Tandem mass spectrometry for the identification and quantification of lipid molecular species was performed using Multiple Reaction Monitoring (MRM) with a TSQ Vantage Triple Stage Quadrupole Mass Spectrometer (Thermo Fisher Scientific) equipped with a robotic nanoflow ion source, Nanomate HD (Advion Biosciences, Ithaca, NY). The collision energy was optimized for each lipid class. The detection conditions for each lipid class are listed in Table S1. Ceramide species were also quantified with a loss of water in the first quadrupole. Each biological replicate was read in 2 technical replicates (TR). Each TR comprised 3 measurements for each transition. Lipid concentrations were calculated relative to the relevant internal standards and then normalized to the total lipid content of each lipid extract (mol%). Data analysis was performed using the LipidSig webtool (Lin et al., 2021) using Benjamini and Hochberg multiple testing correction, adjusted p-value (0.05) and fold change filtering of 1.5.

**Table S1:** Detection of Lipids by MS/MS (Related to Lipidome Analyses). Related to STAR Methods.

| Lipid Class | Standard | Polarity | Mode | m/z ion | Collision Energy |
| --- | --- | --- | --- | --- | --- |
| Phosphatidylcholine [M+H]<sup>+</sup> | DLPC | + | Product ion | 184.07 | 30 |
| Phosphatidylethanolamine [M+H]<sup>+</sup> | PE31:1 | + | Neutral ion loss | 141.02 | 20 |
| Phosphatidylinositol [M-H]<sup>-</sup> | PI31:1 | - | Product ion | 241.01 | 44 |
| Phosphatidylserine [M-H]<sup>-</sup> | PS31:1 | - | Neutral ion loss | 87.03 | 23 |
| Cardiolipin [M-2H]<sup>2-</sup> | CL56:0 | - | Product ion | acyl chain | 32 |
| Ceramide | C17Cer | + | Product ion | 264.34 | 25 |
| Dihydroceramide | C17DHCer | + | Product ion | 266.40 | 25 |
| Hexacylceramide | C8GC | + | Product ion | 264.34 | 30 |
| Hexacyldihydroceramide | C8GC | + | Product ion | 266.40 | 30 |
| Sphingomyelin | C12SM | + | Product ion | 184.07 | 26 |

### UHPLC-HRMS analyses

Dried samples were resuspended by sonicating in 100 µl of LC-MS-grade chloroform:methanol (1:1, v/v). Reversed-phase UHPLC-HRMS analyses were performed using a Q Exactive Plus Hybrid Quadrupole-Orbitrap mass spectrometer coupled to an UltiMate 3000 UHPLC system (Thermo Fisher Scientific) equipped with an Accucore C30 column (150 x 2.1 mm, 2.6 μm) and its 20mm guard column (Thermo Fisher Scientific). Samples were kept at 8°C in the autosampler, 10 μl were injected and eluted with a gradient starting at 10% B for 1 min, 10-70% B in 4 min, 70-100% B in 10 min, washed in 100 % B for 5 min and column equilibration for an additional 3 min. Eluents were made of 5 mM ammonium acetate and 0.1% formic acid in water (*solvent A*) or in isopropanol/acetonitrile (2:1, v/v) (*solvent B*). Flow rate and column oven temperature were respectively at 350 μl/min and 40°C. The mass spectrometer was operated using a heated electrospray-ionization (HESI) source in positive and negative polarity with the following settings: electrospray voltage: −3.4 KV (-) or 3.9 KV (+); sheath gas: 51; auxiliary gas: 13; sweep gas: 3; vaporizer temperature: 431°C; ion transfer capillary temperature: 320 °C; S-lens: 50; resolution: 140,000; m/z range: 200-1000; automatic gain control: 1e6; maximum injection time: 50 ms. For identification of AzoPC and AzoPE, parallel reaction monitoring (PRM) measurement was performed using a predetermined inclusion list of corresponding lipid species. The following setting was used in HCD fragmentation: automatic gain control: 1e6; maximum injection time: 25 ms; resolution: 70,000; (N)CE: 30. Xcaliburv 4.2 (Thermo Fisher Scientific) was used for data acquisition and processing.

### Click Imaging

HeLa cells from an exponentially growing main culture were detached by trypsinization and seeded on a with poly-lysine pre-coated imaging dish in 200 µL medium at a density of 25k cells per well. After the cells attached overnight in the incubator at 37°C and 5 % CO_2_, the medium was carefully removed with a pipette without disturbing the cells. After washing with PBS, HeLa cells were incubated with 200 µL **clFAAzo4** (50 µM in 0.1 % DMSO and PBS) at 37 °C and 5 % CO_2_ for 4 hours. Then, the **clFAAzo4** solution was removed and cells were washed with PBS (1x) before fixation. 200 µL of 4 % paraformaldehyde (in PBS) fixation solution was added to each well and cells were incubated for 20 min at room temperature. Then, the fixation solution was removed and cells were washed with PBS (1x). For the following click-reaction, a master mix was prepared, containing PBS, 5 µM suflo-cyanine-5-azide (from a 5 mM stock in DMSO), 1 mM CuSO_4_ (from a freshly made 20 mM stock in ddH_2_O) and 50 mM sodium ascorbate (from a freshly made 500 mM stock in ddH_2_O). 200 µL of the master mix were added to each well and the cells were incubated in the dark at room temperature for one hour. After the click-reaction, the labeling solution was removed and cells were washed with PBS (1x). Subsequently, a solution of ER painter in PBS was added and cells were incubated at room temperature for 1h and washed again with PBS. Cell imaging was carried out on a Leica DMI6000 B inverted confocal microscope with a Leica HC PL APO 63x/1.30 Glyc CORR CS2 immersion objective to acquire the brightfield and fluorescence images (pinhole 20 µm).

### Epifluorescence Microscopy and Fluorescence Recovery After Photobleaching (FRAP)

For epifluorescence measurements of GUVs, an inverted microscope (IX81, Olympus) was used, which is equipped with a 100x oil immersion objective (NA=1.4, UPlanSApo, Olympus), a mercury short arc (HBO) lamp, and two filter sets for UV-A (λ_exc_=330-385nm, λ_em_≥420nm, P = 0.28 Wcm^-2^) and green (λ_exc_=510-550nm, λ_em_≥590nm, P = 0.35 Wcm^-2^) light illumination. Image sequences were recorded with a CCD camera (iXon, Andor, exposure times: 0.1s). For the FRAP measurements, a small spot on the SLB was photobleached with intense green light for 5 s (excitation filter: 510-550 nm, P = 0.97 W cm^-2^). The fluorescence recovery was recorded by acquiring an image sequence during exposure with UV-A or green light (exposure times: 0.1s). The data were analyzed according to Jönsson *et al*.(Jönsson et al., 2008).

**Preparation of GUVs, SUVs, and SLBs: (i) GUVs** were prepared via electroformation using a home-built device(Urban et al., 2018). Five µl of a mixture of 99 mol% C16-AzoPC and 1 mol% Texas Red-DHPE (ThermoFisher) or 49.5mol% C16-AzoPC, 49.5mol% POPC (Avanti Polar Lipids), and 1 mol% Texas Red-DHPE in CHCl_3_ were spread on the platinum wires of the device using a syringe. The wires (two) are 3 mm apart and span a Teflon chamber. After evaporation of the solvent the chamber was filled with a 300 mM sucrose solution and heated to 70 °C. An actuated electric field (10 Hz, 3 V) was applied to the wires for 120 min. **(ii) SUVs**: 100 µl of lipids dissolved in CHCl_3_ (c = 6.36mM) were added to a glass vial and dried using pressure air. The dry lipids were rehydrated in 1.5 ml ddH_2_O and tip-sonicated (Bandelin Sonoplus) at least two times for 30 s at high intensity until a clear solution was obtained. Finally, the vesicle suspension was centrifuged for 10 min at 8000 rpm. **(iii) SLBs** were made by drop casting SUVs labeled with 1mol% Texas Red-DHPE on glass slides to induce vesicle fusion and bilayer formation.(Lin et al., 2010) The glass substrates were cleaned via sonication in acetone, propanol, and ddH_2_0, each for 5 min, and additional plasma treatment (PDS-32-G, Harrick Plasma) for 1 min. Prior measurement, the samples were rinsed several times to remove excess pSUVs.

### DNA transfection

The amount of DNA per well was 0.05 µg.

Hela MZ cells in 96-well plate*: Trans*IT-X2® Dynamic Delivery System (Mirus Bio) was used to transfect plasmid DNA. Briefly, the medium from each well was replaced with 70 µL of fresh media and a 30 µL DNA solution previously diluted in Opti-MEM and mixed with *Trans*IT-X2® Dynamic Delivery System reagent in a 3:1 *Trans*IT-X2 (µL): DNA (µg) ratio. RUSH constructs were gift from Franck Perez: Str-KDEL_SBP-EGFP-GPI and Str-KDEL_TNF-SBP-mCherry (Addgene plasmid #65294 and #65279).

HCT 116 cells in 96-well plate: DNA transfection was done using Lipofectamine^TM^ 3000 (Cat# L3000008) following manufacturer’s instructions. Briefly, DNA-lipid complexes were prepared in Opti-MEM. Separately, a tube was prepared mixing DNA with P3000 reagent and another tube with Lipofectamine 3000. The content of the tubes was mixed and 30µL of the mixture was added per well to the 70 µL of fresh media previously added.

### RUSH assay

For the high content microscopy experiments we used black 96-well imaging plates (cat# 89626, Ibidi). Plates that were used to seed HCT 116 cells were covered with Poly-D-lysine to improve attachment with the following protocol: 10000 Hela ZM cells or 8000 HCT 116 were seeded per well. Hela MZ cells were transfected with the RUSH plasmids after 24 hours and HCT 116 cells after 48 hours to optimize attachment and as described in the previous section. After 24 hours of DNA transfection, cells were incubated with FAAzo4 as described in previous sections. After 4 hours of treatment, the media in each well was exchanged by fresh media and cells were left in the incubator before the RUSH experiment for at least 15 mins. Next, the 96 well plate was covered with aluminum foil in those wells where the effect of trans-FAAzo4 was to be evaluated and the rest of the wells were illuminated with UV-A light (365 nm) to photoswitch trans-FAAzo4 to cis-FAAzo4. To synchronize and follow the export of GPI-AP or TNF*α* in our cells, we used the approach established by Boncompain *et al*. and previously described in Jiménez-Rojo et al. First, a solution of biotin was prepared in media at a concentration of 120 µM (40 µM final concentration in each well). The cargo (GPI or TNF*α*) construct containing an SBP tag (streptavidin binding protein) and a fluorescent protein (EGFP or mCherry) is fused to minimal ER hook containing streptavidin and a C-terminal ER retention signal (KDEL, Lys-Asp-Glu-Leu) giving rise to the Str-KDEL_SBP-EGFP-GPI or Str-KDEL_TNF-SBP-mCherry construct that is used for transfection. To start the release of the transfected fluorescent cargo, 50 µL of the biotin solution were added at different time points depending on the kinetics of each cargo and on the cell line. Once the biotin solution is added to each well, the reporter is released from the ER hook and follows the secretory pathway. Arrival to the Golgi is measured as explained below, following colocalization of the EGFP-GPI or TNF*α*-mCherry construct with GM130, a Golgi resident protein stained first with a primary purified mouse anti-GM130 antibody (Cat# 610822, BD Biosciences) following staining with a secondary Alexa Fluor® 647-AffiniPure Donkey Anti-Mouse IgG (H+L) (Cat# 715-605-150, Jackson ImmunoResearch) in combination with EGFP-GPI or Alexa Fluor® 488 AffiniPure Donkey Anti-Mouse IgG (H+L) (Cat# 715-545-151, Jackson ImmunoResearch) in combination with TNF*α*-mCherry. After the final time point, the plate was fixed using 4% PFA solution and washed using an automated plate washer (BioTek EL406). Cells were then stained as follows: step 1: monoclonal antibody against GM130, 1/500, saponin 0.05%, BSA 1%, in PBS, incubation for 1 h and, step 2: Hoechst 33342 Solution (20 mM) (1/5000), Cy5 or Alexa 488-labelled secondary antibody against mouse IgG, 1/500, incubation for 30 min. Image acquisition was performed immediately after staining using a ImageXpress® Micro Confocal High-Content Imaging System (Molecular devices) with the 40× objective. 36 images were captured per well. For image analysis, we used the MetaXpress Custom Module editor software to first segment the image and generate relevant masks (Fig S3). In the first step, individual cells were identified using the staining of the nuclei (Hoechst channel). Next, the Golgi was segmented from the images using the signal coming from the anti-GM130 antibody (Cy5 or Alexa 488 channel). Properly transfected cells were then selected using those ones with a fluorescence intensity of the cargo ranging between two specific values, identical through all conditions but different depending on the cargo. Finally, the masks were applied to the original fluorescent images, and different measurements were obtained per cell (e.g. integrated intensity, average intensity and object count). The average intensity value of the fluorescent cargo in the Golgi is used to represent the data. The same imaging and analysis pipelines were applied to all images. Data analysis was with Prism Graph Pad 9.0 and the statistical analysis was performed by unpaired two-tailed Student’s t test with Welch’s correction.

### Analysis of gene transcription

Total RNA from cells was extracted and purified using with RNeasy Mini Kit (Qiagen) and reversely transcribed into cDNA with the iScript cDNA Synthesis Kit (Bio-Rad). Quantitative RT-PCR analysis was performed on a CFX Connect Real-Time PCR system using SsoAdvanced Universal SYBR Green Supermix (Bio-Rad). The primers used were (5’à 3’): GAPDH Foward GGC CAT CCA CAG TCT TCT G GAPDH Reverse TCA TCA GCA ATG CCT CCT G CHOP Foward AGA ACC AGG AAA CGG AAA CAG A CHOP Reverse TCT CCT TCA TGC GCT GCT TT Fold change in transcript levels was calculated using the cycle threshold (CT) comparative method (2^−ddCT^) normalizing to CT values of internal control genes *GAPDH*.

### Generation of ACSL4 mutant cells

ACSL4 polyclonal HeLa MZ mutant cells were generated with CRISPR-Cas9, using the highly efficient co-targeting strategy GENF (Harayama et al., 2020) (GEne co-targeting with Non-eFficient conditions) as following. A plasmid for mammalian cell expression of ACSL4-targeting single-guide RNA (sgRNA) and Cas9 was constructed by annealing the oligonucleotides caccgTCATGGGCTAAATGAATCTG (target sequence in upper case) and aaacCAGATTCATTTAGCCCATGAc and ligating them with Quick Ligase (New England Biolabs) into pX330 plasmid (Addgene #42230, deposited by Feng Zhang) cleaved with FastDigest BpiI (Thermo Scientific). This plasmid (495 ng) was co-transfected with 5 ng of a previously generated mismatched sgRNA expression plasmid (Harayama et al., 2020) to target HPRT1 (target sequence with mismatches in lower cases: gtGCCCTCTGTGTGCTCAA) using Lipofectamine 3000. Five days after transfection, mutant cells were selected with 6 µg/mL 6-thioguanine for one week, which kills cells with a functional HPRT1 gene. 6-thioguanine selected cells having mutated HPRT1 despite the use of mismatched sgRNA and the low amount of plasmid used to express it, enabling the enrichment of cells with high CRISPR-Cas9 activity. The targeted ACSL4 region was amplified from control cells and mutated cells with the PCR primers ACTGATTGCATGCTGTGAATCT and GGTGTGGAGGTCACCAATCAC, the amplicons were treated with Exonuclease I and FastAP Thermosensitive Alkaline Phosphatase (both from Thermo Scientific) and used for Sanger sequencing (Fasteris) with the sequencing primer CAGCTATTAAACTTAAGCCTGC. Mutation efficiency was assessed by analyzing sequence traces with TIDE (Brinkman et al., 2014) (Tracking Indels by DEcomposition). This led to a polyclonal population having undetectable levels of non-mutated ACSL4 alleles.

### Fluorescence Lifetime Imaging Microscopy (FLIM) in Cells

Cells were seeded 3 days before the experiment and treated with 1μM myriocin following the procedure described in Jiménez-Rojo *et al*. and using 35 mm glass-bottom dishes obtained from MatTek (P35G-0.170 14-C). For staining of cells with the ER-flipper to visualize membrane tension, cells were incubated for 20 mins with 1μM ER flipper at 37°C, washed afterwards with live imaging medium (Gibco L-15 Leibovitz’s) and imaged in the same medium with 10% FBS. Fluorescence lifetime imaging microscopy (FLIM) was performed on a Leica SP8 DIVE, at 20 MHz, with *λ*_exc_ = 488 nm (white light laser) and 63x oil immersion objective lens. To measure in ‘inverted mode’, around 200 μL of medium were kept on each dish and a cover slip was placed on top of the surface, then the dish was flipped upside down to image upright without drying out the cells. For analysis, a threshold is applied to exclude background pixels and a biexponential reconvolution model was used for the fitting. ROIs were manually selected from the images, outlining the ER membrane based on the signal of the probe and excluding the plasma membrane which was faintly stained and that could be easily excluded due to higher lifetime values. Each point represents the average lifetime of different ROIs selected from 1 cell.

### Pe-GFP-VSV infection assay

The VSV infection assay was performed using previous published protocols (Le Blanc et al., 2005). We used the same materials and instrument as the RUSH assay unless indicated below. Briefly, 12,000 HeLa cells/well were seeded in 96-well plates using Fluoro-Brite DMEM supplemented with 10% FCS and 2mM L-Glutamine. After 20 h, cells were incubated with 50 µM FAAzo4 or DMSO for 4 h. Next, the 96 well solid black plate was partially covered with aluminum foil for the wells where the effect of trans-FAAzo4 was to be evaluated and remaining wells were illuminated with UV-A light (365 nm) to photoswitch trans-FAAzo4 to cis-FAAzo4. After illumination, cells were washed with cold PBS (3x), and supplemented with 100 µL cold VSVmem (Glasgow Minimum Essential Medium, 10mM TES, 10mM MOPS, 15mM HEPES, 2mM NaH2PO4, 35 mg/L NaHCO3, pH = 7.4), incubated for 5 min on ice. Cells were then treated with 50 µL of Pe-GFP-VSV virus to reach a final concentration of 0.5 MOI, incubated for 45 min on ice with gentle shaking, washed by PBS (3x), supplemented with 150 µL of growth medium (37°C), incubated for another 3.5 h before fixation using 3% paraformaldehyde. For staining, the plate was washed (3x) by PBS, cells were incubated with Hoechst in PBS for 30 min and washed again (3x). Image acquisition was performed immediately after staining using a ImageXpress® Micro Confocal High-Content Imaging System (Molecular devices) with the 20× objective. For image analysis, we used the MetaXpress Custom Module editor software to first segment the image and generate relevant masks. Cells were scored as infected by the presence of the Pe-GFP-VSV protein by automated image analysis.

### Phophoproteomics

#### Cell lysis and protein digestion

Cell pellets were suspended in lysis buffer composed of 8M urea 100 mM TRIS pH=8.5 and lysed by sonication. Protein concentrations were measured by BCA protein assay. Lysates were supplemented with TCEP (final 5 mM), chloroacetamide (final 10 mM) and incubated @ 56 °C for 1h. After 6X dilution with 25 mM ammonium bicarbonate, proteins were digested with trypsin (100:1 ratio, o/n @ 37°C). Digestion was stopped by acidification with FA (final 0.5%) and peptides were desalted on tC18 cartridges (50 mg, 1cc, Waters) according to manufacturer instructions. Peptide elution was performed in 400 ul of 40% ACN and 200 ul of 60% ACN w/o any acid. Peptide concentrations were measured by Pierce colorimetric peptide assay, subsequently these measurements were used for TMT labeling. Finally, all samples were dried in speedvac and stored at −80°C.

#### TMT labeling

Each peptide sample were reconstituted in 20 ul of 50 mM HEPES, pH=8.5. TMT labeling was performed with TMTPro isobaric tags according to procedure adapted from(Zecha et al., 2019). Briefly, samples were labeled with 8 ul of the corresponding TMT label ACN stock (12.5 ug/ul of label). Samples were incubated at RT for 30min before quenching with 40 ul of 500 mM ABC (15 min at 37°C). To lower ACN concentration prior to the desalting step each sample was diluted with an additional 400 ul of 0.5% TFA. All TMT channels were pooled together and desalted on tC18 cartridge (50 mg, 1cc, Waters) according to manufacturer instructions. Small aliquots were used as a QC for labeling completion. Eluates were dried in a speedvac concentrator and stored at −80°C.

#### Phosphopeptide enrichment

Phosphorylated peptides were enriched by IMAC on high-select Fe-NTA spin columns (Thermo Scientific) according to the manufacturer instructions. Eluted pSTY peptides were dried in a speedvac concentrator and resolubilized in 10 ul of 2% ACN 0.5% AcOH prior to LC-MS/MS analysis. **LC-MS/MS.** LC separation was performed online on EASY-nLC 1000 (Thermo Scientific) utilizing Acclaim PepMap 100 (75 um x 2 cm) precolumn and PepMap RSLC C18 (2 um, 100A x 50 cm) analytical column. Peptides were gradient eluted from the column directly into an Orbitrap HFX mass spectrometer using 136 min ACN gradient from 5 to 26 % B in 100 min followed by ramp to 40% B in 20 min and final equilibration in 100% B for 15 min (A=2% ACN 0.5% AcOH / B=80% ACN 0.5% AcOH). Flowrate was set at 200 nl/min. High resolution full MS spectra were acquired with a resolution of 120,000, an AGC target of 3e6, with a maximum ion injection time of 100 ms, and scan range of 400 to 1600 m/z. Following each full MS scan 20 data-dependent HCD MS/MS scans were acquired at a resolution of 60,000, AGC target of 5e5, maximum ion time of 100 ms, one microscan, 0.4 m/z isolation window, NCE of 30, fixed first mass 100 m/z and dynamic exclusion for 45 seconds. Both MS and MS^2^ spectra were recorded in profile mode.

## Data Analysis

MS data were analyzed using MaxQuant software version 1.6.3.4(Cox and Mann, 2008) and searched against the SwissProt subset of the human uniprot database (http://www.uniprot.org/) containing 20,430 entries. Database search was performed in Andromeda(Cox et al., 2011a) integrated in MaxQuant environment. A list of 248 common laboratory contaminants included in MaxQuant was also added to the database as well as reversed versions of all sequences. For searching, the enzyme specificity was set to trypsin with the maximum number of missed cleavages set to 2. The precursor mass tolerance was set to 20 ppm for the first search used for non-linear mass re-calibration(Cox et al., 2011b) and then to 6 ppm for the main search. Phosphorylation of S/T/Y and oxidation of methionine were searched as variable modification; carbamidomethylation of cysteines was searched as a fixed modification. TMT labeling was set to lysine residues and N-terminal amino groups, corresponding batch-specific isotopic correction factors were accounted for. The false discovery rate (FDR) for peptide, protein, and site identification was set to 1%, the minimum peptide length was set to 6. Subsequent data analysis were performed in either Perseus(Tyanova et al., 2016) (http://www.perseus-framework.org/) or using R environment for statistical computing and graphics (http://www.r-project.org/).

## Supporting Information

^1^H and ^13^C NMR spectra and Supplementary Figures.

## Author Contributions

S.F., J.M., H.R., and D.T. conceived the study. N.J.R., S.F., and J.M. performed most experiments and data analysis. S.D.P., A.A., N.A.V., C.J.A., M.R., E.K., B.U., T.L. performed experiments and data analysis. T.H. and A.J.E.N. provided unpublished reagents. N.J.R., S.F., J.M., H.R., and D.T. wrote the manuscript with input from all authors.

## Acknowledgments

J.M. thanks the NCI for a K00 award (K00CA253758). J.M. and N.A.V. thank the German Academic Scholarship Foundation (Studienstiftung des deutschen Volkes) for a PhD fellowship. J.M. thanks New York University for a MacCracken and Margaret and Herman Sokol Fellowship. This work was supported by grants from the Swiss National Science Foundation and the National Centre for Competence in Research in Chemical Biology (310030-184949, 51NF40-185898 to HR) and from the Leducq Foundation (HR) and from the Deutsche Forschungsgemeinschaft (DFG, German Research Foundation) – Project-ID 201269156 – SFB 1032. We would like to thank the laboratory of Prof. Stefan Matile in the University of Geneva for sharing the ER-specific mechanosensitive probe. The authors thank Damien Marechal and Patrizia Casaccia for additional uptake studies (not included). We would like to thank Dimitri Moreau and Stefania Vossio from ACCESS Geneva (University of Geneva) for technical support on the high content microscopy experiments.

---

**Figure S1.**
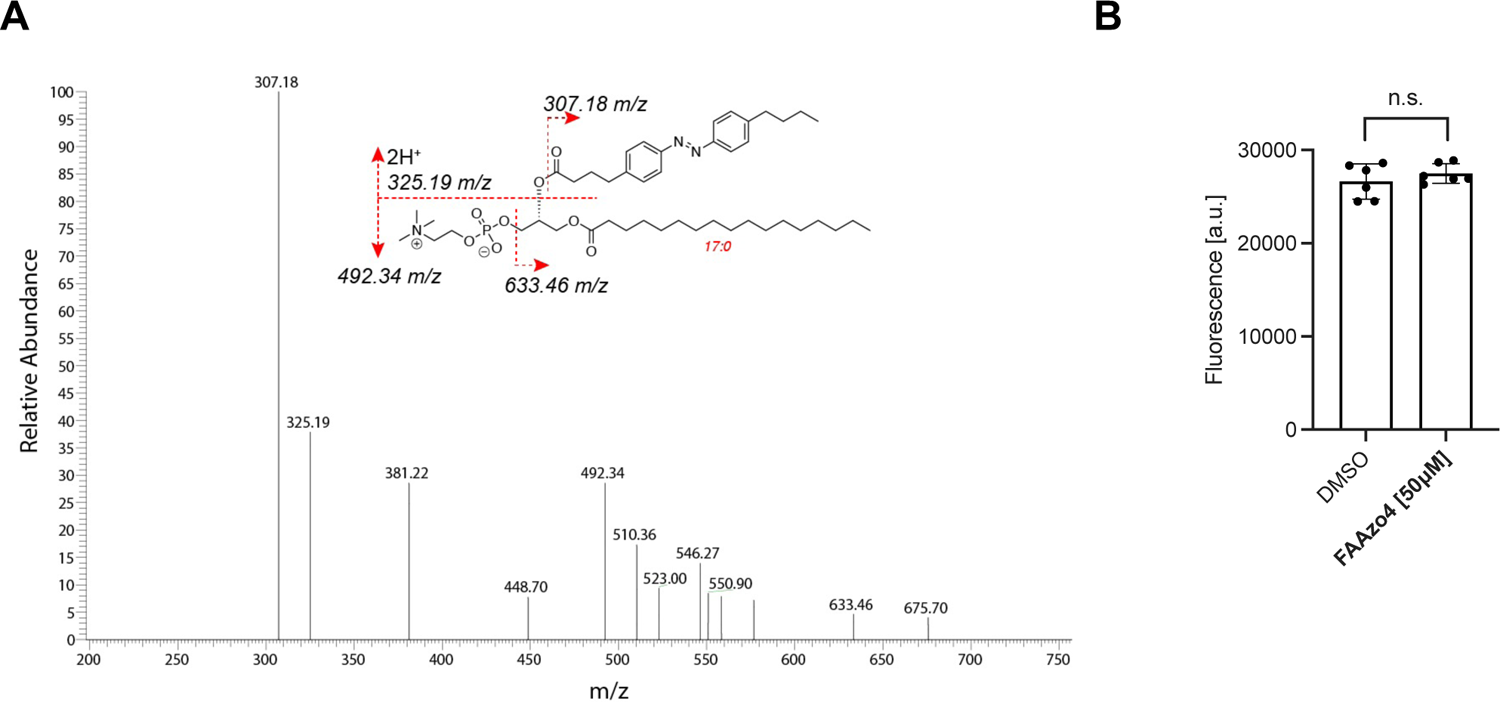
LCMS characterization of C17-AzoPC and cell viability of HeLa PhotoCells. (A) LC-MS/MS spectra of **C17-AzoPC**. (B) Cell viability of Hela cells after incubation with FAAzo4 (50 μM) for 4h.

**Figure S2.**
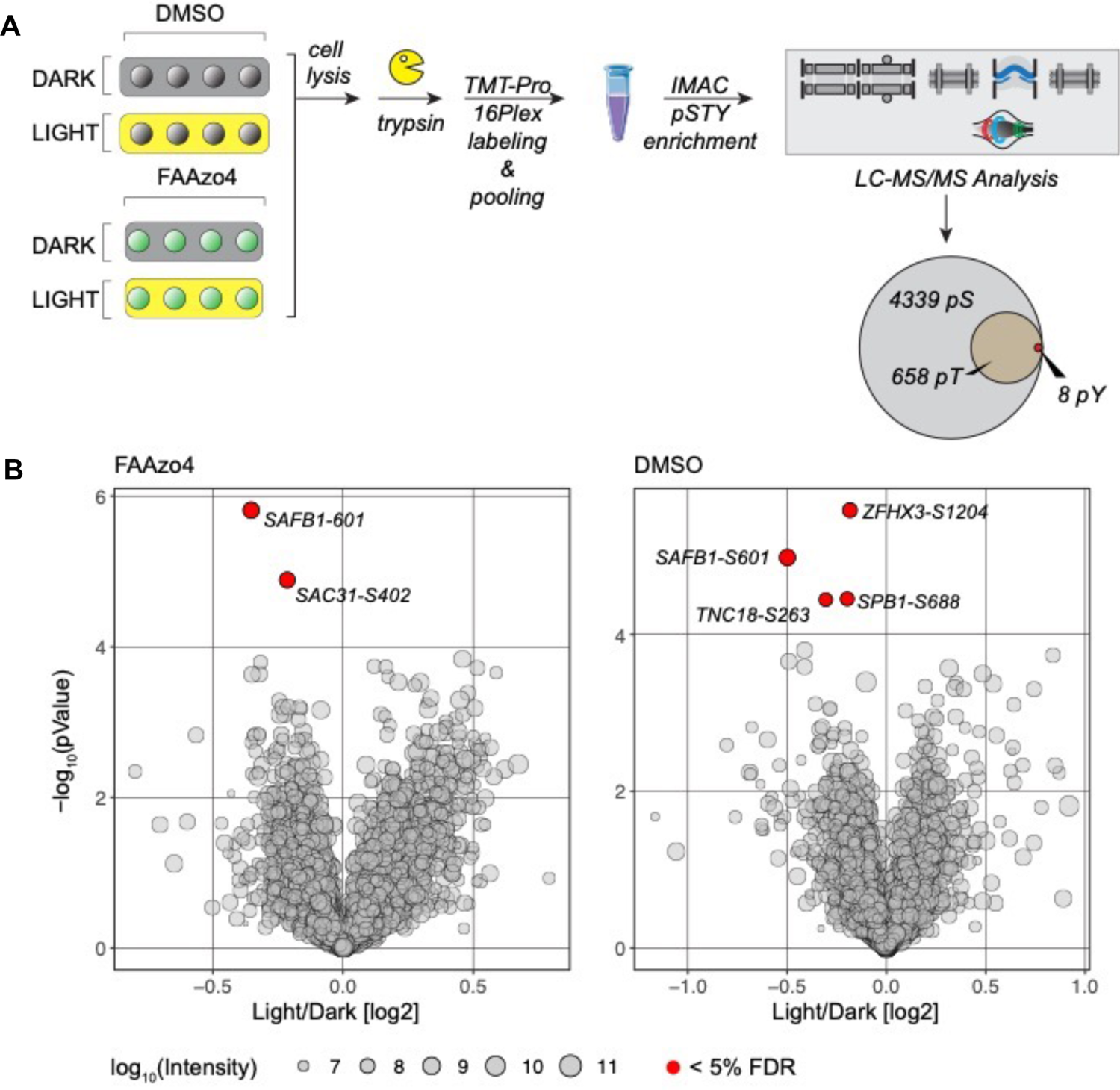
Light-dependent phosphoproteomics in HeLa photocells. (A) **FAAzo4** and control HeLa cell pellets from both dark/light-stimulated conditions (n=4 from each group) were processed in parallel. Cells were lysed in urea buffer and proteins were digested into peptides with trypsin. Peptides from each sample were labeled with TMT-Pro isobaric tags and all samples were pooled together. Phosphopeptides were purified by IMAC and subsequently analyzed by LC-MS/MS, which resulted in identification and quantification of ∼ 5,000 phosphosites. (B) Statistical analyses revealed only a minor difference between light/dark conditions (permutation-based FDR threshold of 5%). We did not observe more difference within **FAAzo4**-trated cells compared to controls.

**Figure S3.**
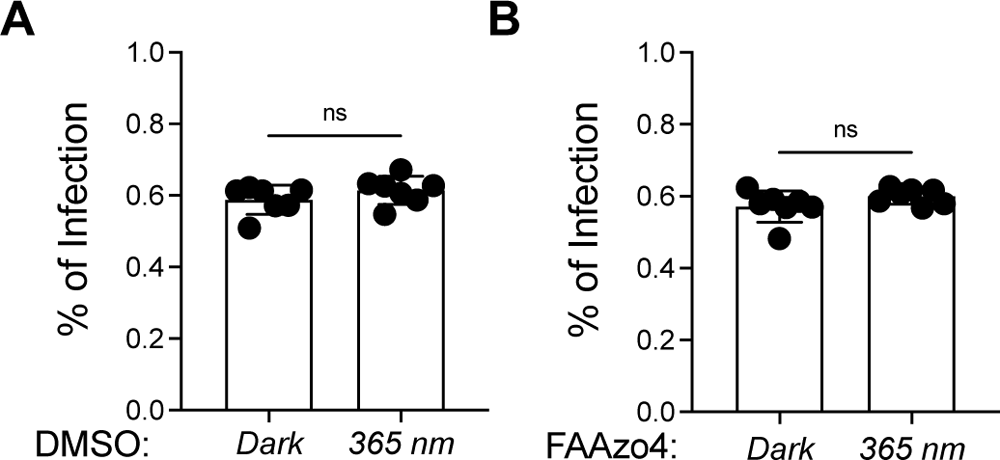
Light-dependent VSV infection assay in HeLa photocells. FAAzo4. (A) or DMSO (B) treated cells were illuminated with UV-A light at 365 nm and control wells were covered with aluminum foil. After the infection experiments, cells were fixed, stained with Hoechst, and images were acquired using automatic fluorescence microscopy.

**Figure S4.**
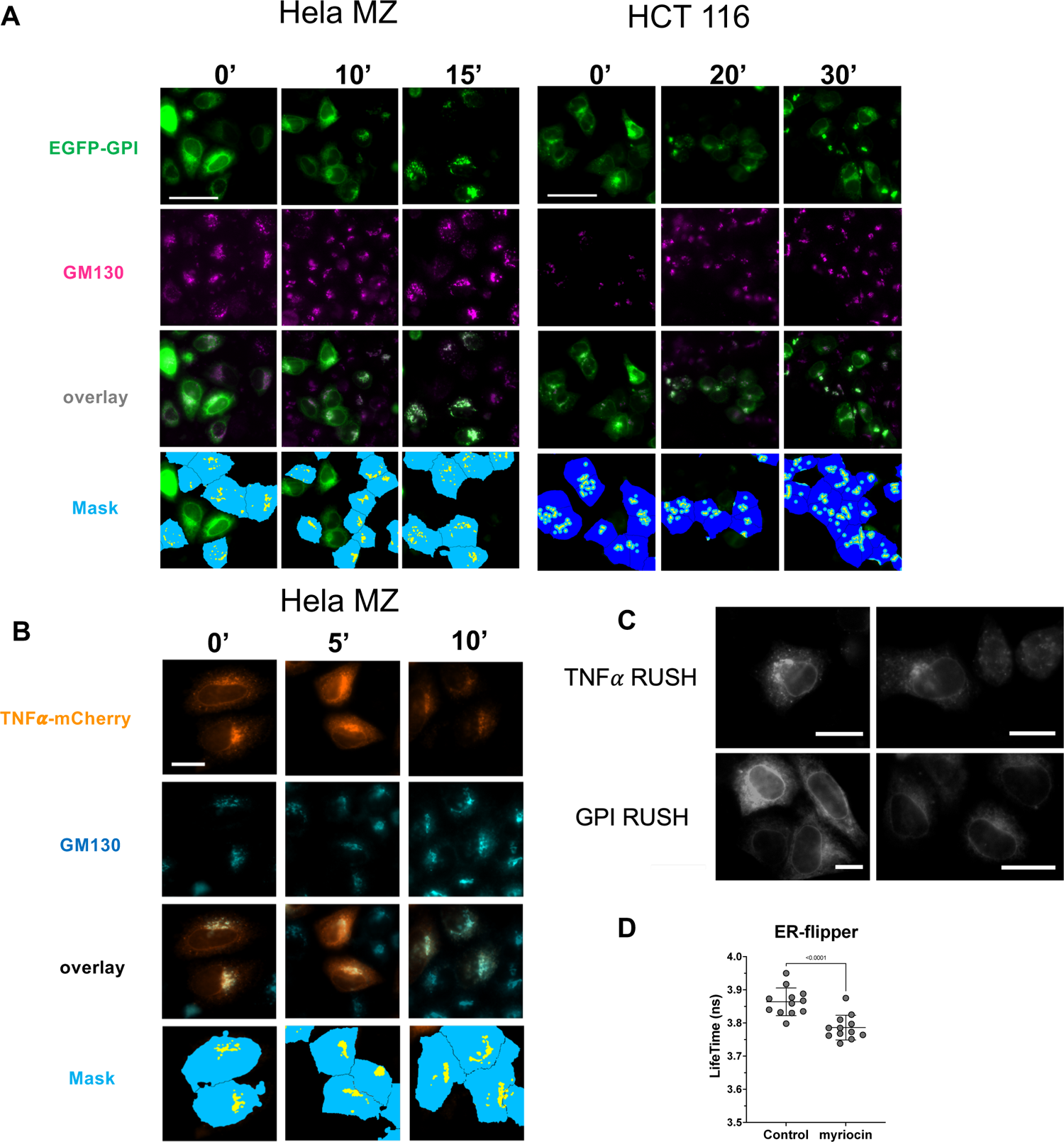
Representative images of the RUSH assay in different cell lines. Related to. Figure 5**. (A)** Representative microscopy images of the RUSH experiment following GPI-RUSH in HeLa cells and HCT 116 cells. Time points represent the time after biotin addition. (B) Representative microscopy images of the RUSH experiment following TNF*α*-RUSH in HeLa cells. Time points represent the time after biotin addition. (C) Representative images showing expressed selected cargo in HeLa cells before release upon biotin addition. (D) Fluorescence lifetime values of the ER-flipper probe in HeLa cells treated with myriocin.

**Figure S5.**
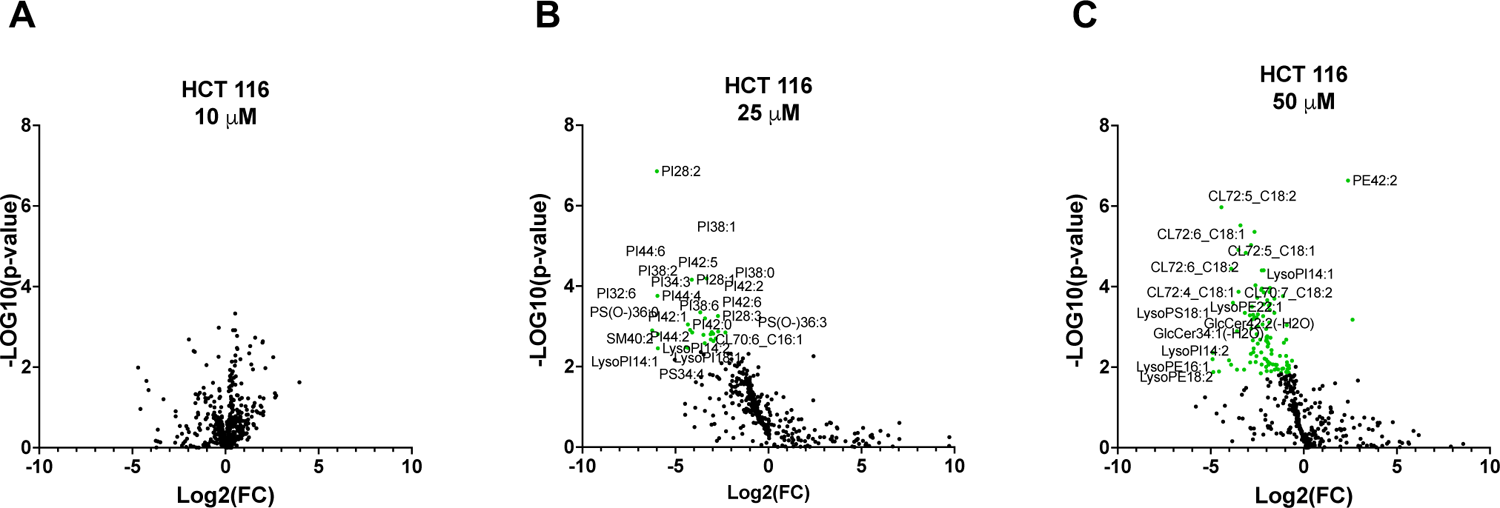
Lipidomic analysis of HCT cells treated with increasing amounts of FAAzo4. Related to. Figure 6. Data and statistical analysis were obtained using LipiSig Webtool. (Lin et al., 2021) (A) 10 μM. (B) 25 μM. (C) 50 μM.

## Synthetic Scheme

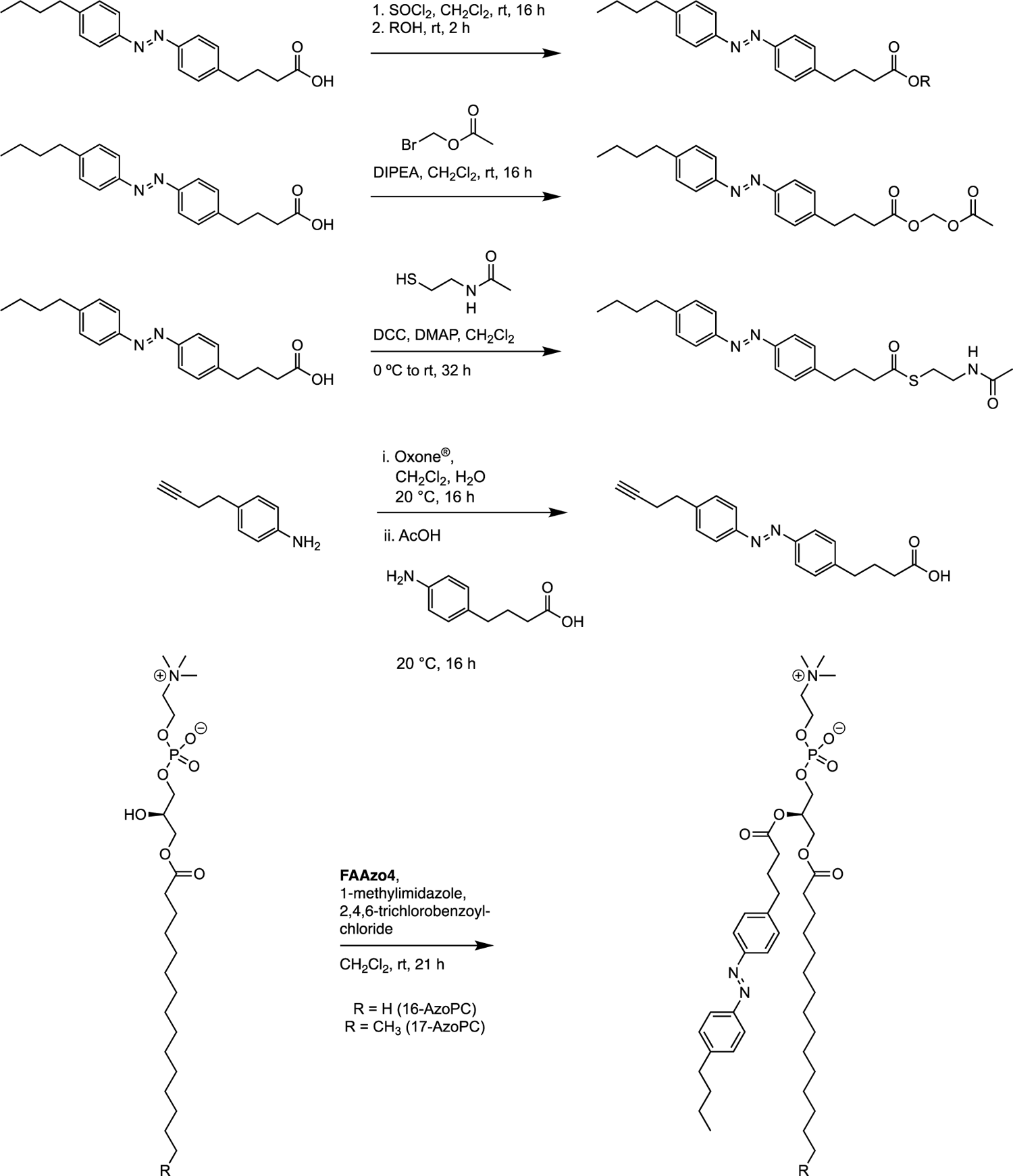

### FAAzo4-Me

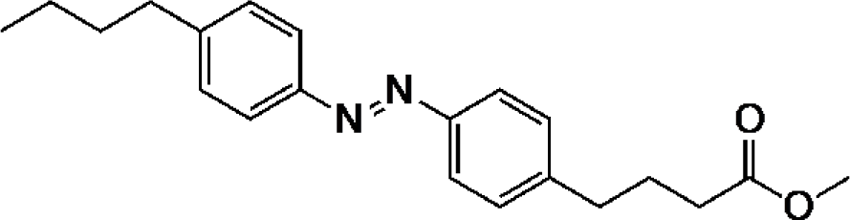

FAAzo4^1^ (30.0 mg, 92.5 μmol, 1.00 equiv.) was dissolved in a 1 M SOCl_2_ (10.0 mL, 10.0 mmol, 108 equiv.) solution in CH_2_Cl_2_ and stirred at rt for 16 h. Volatiles were removed under reduced pressure, residues were dissolved in 1.3 mL CH_2_Cl_2_, and MeOH (37.0 μL, 0.925 mmol, 10.0 equiv.) was added. The mixture was stirred at rt for 2 h. The solvent was removed under reduced pressure. The crude was purified using flash column chromatography (hexanes to 30 % EtOAc in hexanes) to yield the product (24.5 mg, 72.4 μmol, 78%) as an orange liquid.

**^1^H NMR** (400 MHz, CDCl_3_) δ 7.83 (dd, *J* = 8.4, 2.3 Hz, 4H), 7.31 (d, *J* = 8.1 Hz, 4H), 3.68 (s, 3H), 2.77 – 2.65 (m, 4H), 2.36 (t, *J* = 7.4 Hz, 2H), 2.01 (p, *J* = 7.5 Hz, 2H), 1.65 (p, *J* = 7.5 Hz, 2H), 1.38 (h, *J* = 7.3 Hz, 2H), 0.94 (t, *J* = 7.3 Hz, 3H).

**^13^C NMR** (100 MHz, CDCl_3_) δ 174.0, 151.4, 151.1, 146.5, 144.7, 129.3, 129.2, 123.0, 122.9, 51.7, 35.7, 35.1, 33.6, 33.5, 26.5, 22.5, 14.1.

**HRMS**: m/z calc. for C_21_H_27_N_2_O_2_^+^ ([M+H]^+^): 339.2067, found: 339.2073.

### FAAzo4-Et

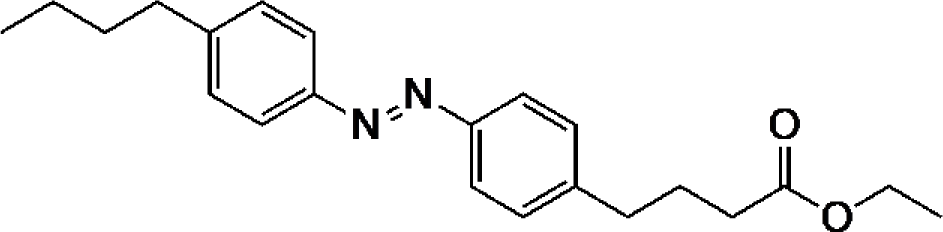

FAAzo4^1^ (30.0 mg, 92.5 μmol, 1.00 equiv.) was dissolved in a 1 M SOCl_2_ (10.0 mL, 10.0 mmol, 108 equiv.) solution in CH_2_Cl_2_ and stirred at rt for 16 h. Volatiles were removed under reduced pressure, residues were dissolved in 1.3 mL CH_2_Cl_2_, and EtOH (54.0 μL, 0.925 mmol, 10.0 equiv.) was added. The mixture was stirred at rt for 2 h. The solvent was removed under reduced pressure. The crude was purified using flash column chromatography (hexanes to 30 % EtOAc in hexanes) to yield the product (30.4 mg, 86.2 μmol, 93%) as an orange liquid.

**^1^H NMR** (400 MHz, CDCl_3_) δ 7.83 (dd, *J* = 8.4, 2.3 Hz, 4H), 7.31 (d, *J* = 7.0 Hz, 4H), 4.20 – 4.10 (m, 2H), 2.71 (dt, *J* = 18.3, 7.6 Hz, 4H), 2.35 (t, *J* = 7.4 Hz, 2H), 2.00 (p, *J* = 7.5 Hz, 2H), 1.66 (q, *J* = 7.5 Hz, 2H), 1.38 (h, *J* = 7.3 Hz, 2H), 1.26 (t, *J* = 7.1 Hz, 3H), 0.94 (t, *J* = 7.3 Hz, 3H).

**^13^C NMR** (100 MHz, CDCl_3_) δ 173.5, 151.4, 151.1, 146.5, 144.7, 129.3, 129.2, 123.0, 122.9, 60.5, 35.7, 35.2, 33.7, 33.6, 26.5, 22.5, 14.4, 14.1.

**HRMS**: m/z calc. for C_22_H_29_N_2_O_2_^+^ ([M+H]^+^): 353.2224, found: 353.2228.

### FAAzo4-Bu

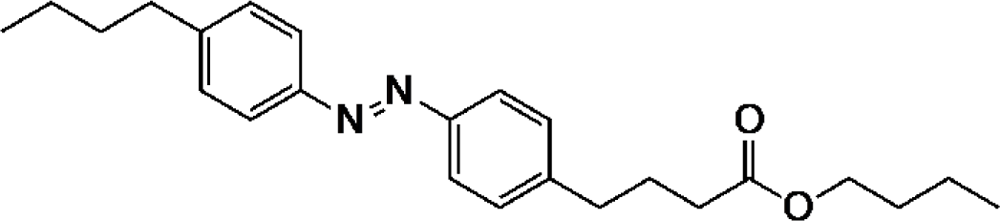

FAAzo4^1^ (30.0 mg, 92.5 μmol, 1.00 equiv.) was dissolved in a 1 M SOCl_2_ (10.0 mL, 10.0 mmol, 108 equiv.) solution in CH_2_Cl_2_ and stirred at rt for 16 h. Volatiles were removed under reduced pressure, residues were dissolved in 1.3 mL CH_2_Cl_2_, and BuOH (85.0 μL, 0.925 mmol, 10.0 equiv.) was added. The mixture was stirred at rt for 2 h. The solvent was removed under reduced pressure. The crude was purified using flash column chromatography (hexanes to 30 % EtOAc in hexanes) to yield the product (28.8 mg, 75.7 μmol, 82%) as an orange liquid.

**^1^H NMR** (400 MHz, CDCl_3_) δ 7.83 (dd, *J* = 8.4, 2.4 Hz, 4H), 7.31 (d, *J* = 8.0 Hz, 4H), 4.08 (t, *J* = 6.7 Hz, 2H), 2.71 (dt, *J* = 17.8, 7.6 Hz, 4H), 2.35 (t, *J* = 7.4 Hz, 2H), 2.00 (p, *J* = 7.5 Hz, 2H), 1.63 (tq, *J* = 12.9, 7.0 Hz, 4H), 1.38 (h, *J* = 7.4 Hz, 4H), 0.94 (td, *J* = 7.3, 3.1 Hz, 6H).

**^13^C NMR** (100 MHz, CDCl_3_) δ 173.6, 151.4, 151.1, 146.5, 144.7, 129.3, 129.2, 123.0, 122.9, 64.4, 35.7, 35.2, 33.8, 30.8, 26.5, 22.5, 19.3, 14.1, 13.9.

**HRMS**: m/z calc. for C_24_H_33_N_2_O_2_^+^ ([M+H]^+^): 381.2537, found: 381.2534.

### FAAzo4-Am

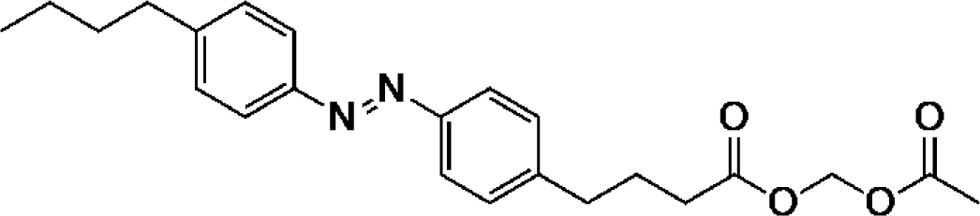

FAAzo4^1^ (50.0 mg, 0.154 mmol, 1.00 equiv.) was dissolved in 2 mL CH_2_Cl_2_, DIPEA (48.0 μL, 0.277 mmol, 1.80 equiv.), and acetoxymethyl bromide (47.2 mg, 0.308 mmol, 2.00 equiv.) were added, and stirred at rt for 16 h. Volatiles were removed under reduced pressure and the crude prduct was purified using flash column chromatography (hexanes to 30 % EtOAc in hexanes) to yield the product (12.5 mg, 31.5 μmol, 20%) as an orange liquid.

**^1^H NMR** (400 MHz, CDCl_3_) δ 7.83 (dd, *J* = 8.3, 2.5 Hz, 4H), 7.31 (dd, *J* = 8.3, 1.6 Hz, 4H), 5.75 (s, 2H), 2.71 (dt, *J* = 20.6, 7.6 Hz, 4H), 2.41 (t, *J* = 7.4 Hz, 2H), 2.03 (q, *J* = 7.5 Hz, 2H), 1.65 (p, *J* = 7.6 Hz, 2H), 1.38 (dq, *J* = 14.7, 7.4 Hz, 2H), 0.94 (t, *J* = 7.3 Hz, 3H).

**^13^C NMR** (100 MHz, CDCl_3_) δ 172.2, 169.8, 151.5, 151.1, 146.5, 144.3, 129.3, 129.2, 123.0, 122.9, 79.3, 35.7, 34.9, 33.6, 33.3, 26.1, 22.5, 20.9, 14.1.

**HRMS**: m/z calc. for C_23_H_29_N_2_O_4_^+^ ([M+H]^+^): 397.2122, found: 397.2123.

### FAAzo4-SNAC

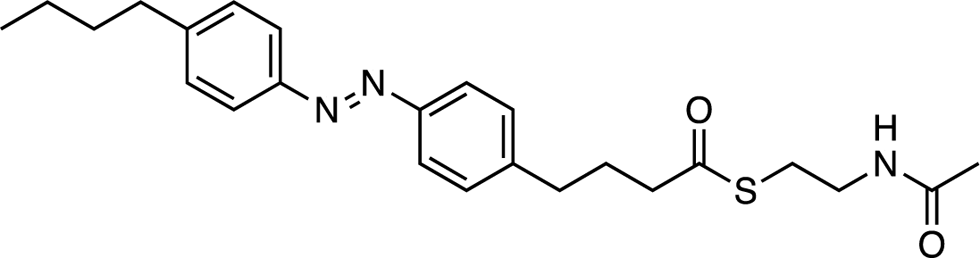

FAAzo4^1^ (50.0 mg, 0.154 mmol, 1.00 equiv.) was dissolved in 2 mL CH_2_Cl_2_ at 0 °C. A solution of DCC (35.0 mg, 0.170 mmol, 1.10 equiv) and DMAP (1.9 mg, 0.015 mmol, 0.10 equiv.) in 1 mL CH_2_Cl_2_ were added. 2-(acetylamino)ethanethiol (19.3 mg, 0.162 mmol, 1.05 equiv.) was added at 0 °C and stirred at rt for 32 h. Volatiles were removed under reduced pressure and the crude prduct was purified using flash column chromatography (15 % EtOAc in hexanes to 80 % EtOAc in hexanes) to yield the product (60.8 mg, 0.143 mmol, 93%) as an orange solid.

**^1^H NMR** (400 MHz, CDCl_3_) δ 7.82 (dd, *J* = 8.4, 3.4 Hz, 4H), 7.36 – 7.29 (m, 4H), 3.42 (q, *J* = 6.2 Hz, 2H), 3.03 (t, *J* = 6.4 Hz, 2H), 2.77 – 2.66 (m, 4H), 2.61 (t, *J* = 7.4 Hz, 2H), 2.05 (q, *J* = 8.6, 8.1 Hz, 2H), 1.95 (s, 3H), 1.69 – 1.62 (m, 2H), 1.39 (dt, *J* = 14.9, 7.4 Hz, 2H), 0.94 (t, *J* = 7.3 Hz, 3H).

**^13^C NMR** (100 MHz, CDCl_3_) δ 199.7, 170.4, 151.5, 151.1, 146.5, 144.2, 129.3, 129.2, 123.0, 122.9, 43.3, 39.8, 35.7, 34.9, 33.6, 28.7, 27.0, 23.4, 22.5, 14.1.

**HRMS**: m/z calc. for C_24_H_32_N_3_O_2_S^+^ ([M+H]^+^): 426.2210, found: 426.2209.

### clFAAzo4

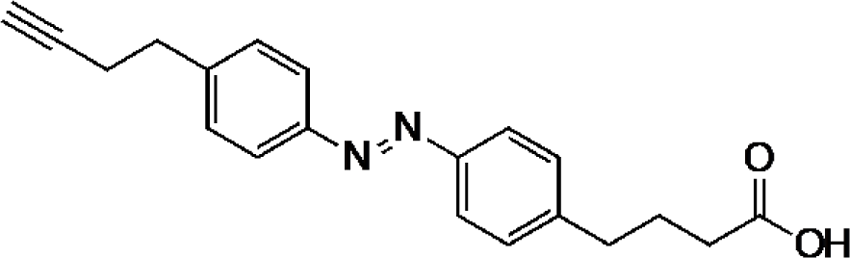

4-(but-3-yn-1-yl)aniline (22.0 mg, 0.123 mmol, 1.00 equiv.) was dissolved in 4 mL CH_2_Cl_2_. A solution of Oxone^®^ (340 mg, 0.552 mmol, 4.50 equiv.) in 4 mL H_2_O was added and stirred vigorously at rt for 16 h. The organic phase was washed with 1 M HCl, a saturated solution of NaHCO_3_, water, dried over Na_2_SO_4_, and filtered. 4-(4-aminophenyl)butyric acid (26.7 mg, 0.184 mmol, 1.50 equiv.) and acetic acid (4 mL) were added, CH_2_Cl_2_ was removed under reduced pressure, and the solution was stirred for 16 h at room temperature. Acetic acid was removed under reduced pressure, azeotroped with toluene, and the product was purified by flash column chromatography (1% AcOH in CH_2_Cl_2_) to yield **clFAAzo4** (16.5 mg, 51.5 μmol, 42%) as an orange solid.

**^1^H NMR** (400 MHz, CDCl_3_) δ 7.88 – 7.81 (m, 4H), 7.35 (dd, *J* = 18.3, 8.4 Hz, 4H), 2.93 (t, *J* = 7.4 Hz, 2H), 2.75 (h, *J* = 7.4 Hz, 2H), 2.54 (td, *J* = 7.4, 2.6 Hz, 2H), 2.41 (t, *J* = 7.4 Hz, 2H), 2.06 – 1.94 (m, 3H).

**^13^C NMR** (100 MHz, CDCl_3_) δ 178.3, 151.6, 151.4, 144.6, 143.6, 129.3, 129.3, 123.1, 123.0, 83.5, 69.4, 35.0, 34.8, 33.1, 26.2, 20.5.

**HRMS**: m/z calc. for C_20_H_21_N_2_O_2_^+^ ([M+H]^+^): 321.1598, found: 321.1594.

### 16-AzoPC (16:FAAzo4)

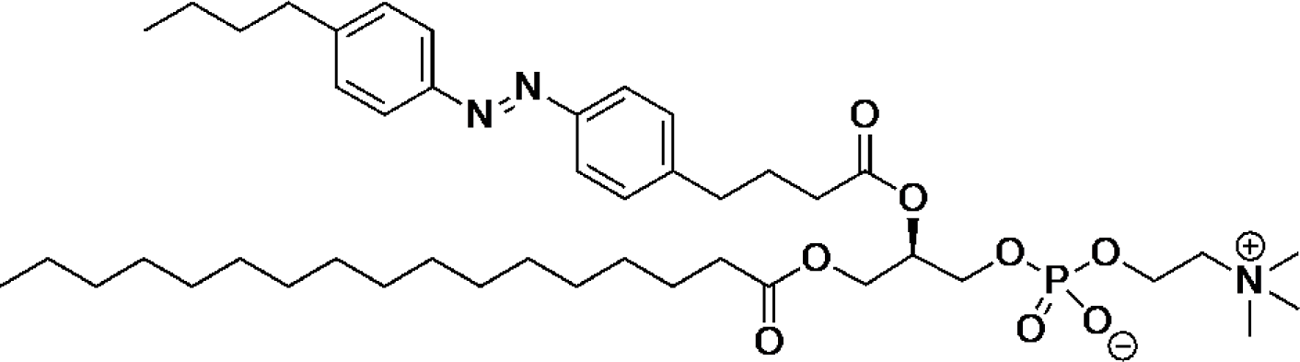

16-LysoPC (50.0 mg, 0.101 mmol, 1.00 equiv.) was dissolved in 30 mL of CH_2_Cl_2_. 2,4,6-trichlorobenzoyl chloride (73.8 mg, 0.302 mmol, 3.0 equiv.), **FAAzo-4**^1^ (65.5 mg, 202 mmol, 2.0 equiv.), and NMI (*N*-methylimidazol) (24.8 mg, 0.302 mmol, 3.0 equiv.) were added to the solution. The clear orange solution was stirred for 16 h at room temperature and was directly subjected to purification via flash column chromatography (CH_2_Cl_2_, CH_2_Cl_2_:MeOH:H_2_O 99:1:0 / 95:5:0 / 8:2:0,1 / 7.5:2.5:0.1 / 7:3:0,1 / 7:3:0,2 / 10:4:0.5) to yield **16-AzoPC (16:FAAzo4)** (15.6 mg, 19.5 μmol, 19 %) as an orange glue.

**^1^H NMR** (400 MHz, MeOD) δ 7.83 (t, *J* = 8.1 Hz, 4H), 7.37 (dd, *J* = 13.1, 8.4 Hz, 4H), 5.32 – 5.22 (m, 1H), 4.50 – 4.41 (m, 1H), 4.28 (d, *J* = 4.5 Hz, 2H), 4.25 – 4.15 (m, 1H), 4.02 (q, *J* = 6.1 Hz, 2H), 3.63 (dd, *J* = 5.6, 3.3 Hz, 2H), 3.21 (s, 9H), 2.74 (dt, *J* = 20.8, 7.7 Hz, 4H), 2.47 – 2.36 (m, 2H), 2.34 – 2.27 (m, 2H), 2.00 (dq, *J* = 11.8, 6.8 Hz, 2H), 1.71 – 1.62 (m, 2H), 1.56 (p, *J* = 7.4 Hz, 2H), 1.42 (dd, *J* = 15.0, 7.5 Hz, 2H), 1.31 – 1.19 (m, 32H), 0.97 (t, *J* = 7.4 Hz, 3H), 0.89 (t, *J* = 6.8 Hz, 4H).

**^13^C NMR** (100 MHz, MeOD) δ 175.0, 174.2, 152.5, 152.3, 147.8, 146.4, 130.4, 130.2, 123.9, 123.8, 72.1, 64.9, 63.6, 60.5, 60.5, 54.7, 54.7, 54.6, 49.7, 49.6, 49.5, 49.4, 49.3, 49.2, 49.1, 49.0, 48.8, 48.6, 48.4, 36.5, 35.8, 34.9, 34.8, 34.4, 33.1, 30.8, 30.6, 30.5, 30.4, 30.2, 27.7, 26.0, 23.7, 23.4, 14.5, 14.3.

**^31^P NMR** (162 MHz, MeOD) δ −0.6.

### 17-AzoPC (17:FAAzo4)

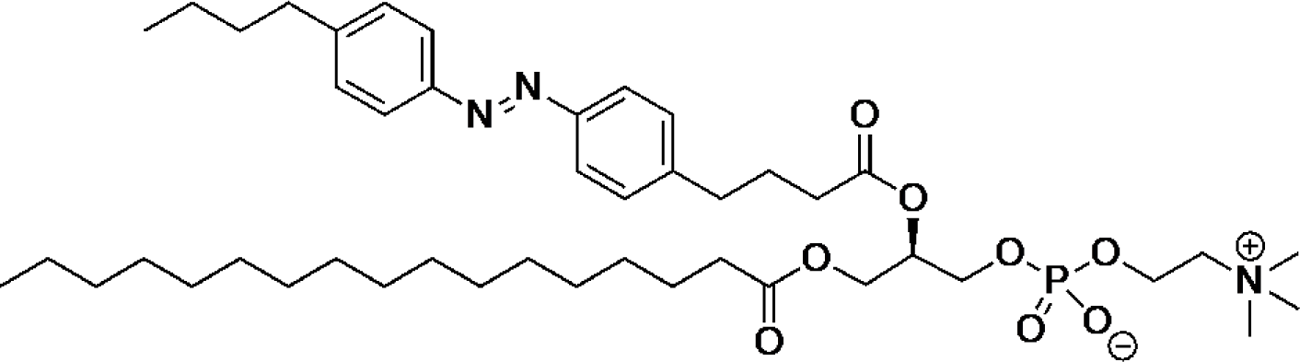

17-LysoPC (10.0 mg, 19.6 μmol, 1.0 equiv.) was dissolved in 620 μL of CH_2_Cl_2_. 2,4,6-trichlorobenzoyl chloride (14.4 mg, 58.9 μmol, 3.0 equiv.), **FAAzo-4**^1^ (12.7 mg, 39.2 μmol, 2.0 equiv.), and NMI (*N*-methylimidazol) (4.8 mg, 58.9 μmol, 3.0 equiv.) were added to the solution. The clear orange solution was stirred for 21 h at room temperature and was directly subjected to purification via flash column chromatography (CH_2_Cl_2_, CH_2_Cl_2_:MeOH:H_2_O 99:1:0 / 95:5:0 / 8:2:0,1 / 7.5:2.5:0.1 / 7:3:0,1 / 7:3:0,2 / 10:4:0.5) to yield **17-AzoPC (17:FAAzo4)** (15.9 mg, 19.5 μmol, 99 %) as an orange glue.

**^1^H NMR** (400 MHz, MeOD) δ 7.83 (t, *J* = 8.0 Hz, 4H), 7.42 – 7.33 (m, 4H), 5.28 (s, 1H), 4.49 – 4.43 (m, 1H), 4.27 (s, 2H), 4.20 (dd, *J* = 12.0, 6.8 Hz, 1H), 4.05 – 3.96 (m, 2H), 3.62 (s, 2H), 3.21 (s, 9H), 2.79 – 2.67 (m, 4H), 2.41 (t, *J* = 7.3 Hz, 2H), 2.30 (t, *J* = 7.5 Hz, 2H), 2.04 – 1.95 (m, 2H), 1.65 (q, *J* = 7.8 Hz, 2H), 1.56 (p, *J* = 7.2 Hz, 2H), 1.41 (dt, *J* = 14.9, 7.4 Hz, 2H), 1.21 (s, 31H), 0.97 (t, *J* = 7.4 Hz, 3H), 0.89 (t, *J* = 6.7 Hz, 3H).

**^13^C NMR** (100 MHz, MeOD) δ 173.5, 172.8, 151.1, 150.9, 146.4, 145.0, 129.0, 128.8, 122.5, 122.4, 70.7, 70.6, 66.1, 63.5, 62.2, 59.1, 59.0, 53.3, 53.2, 35.1, 34.4, 33.5, 33.4, 33.0, 31.7, 29.4, 29.4, 29.2, 29.1, 29.0, 28.8, 26.3, 24.6, 22.3, 22.0, 13.1, 12.9.

**^31^P NMR** (162 MHz, MeOD) δ −0.6.

